# Scientific exploration outlasts industrial momentum and sustains innovation in genetic research: survey of 20 million papers and patents

**DOI:** 10.1101/2025.02.17.638429

**Authors:** Junghun Chae, WooJoong Kim, Woochul Jung, Dawoon Jeong, Roktaek Lim, Manoj Chamlagain, Giju Jung, Juneil Jang, Jae Won Lee, Nam Kyu Kang, Kwangryul Baek, Jonghyeok Shin, Ye-Gi Lee, Hyun Gi Koh, Chanwoo Kim, Sangdo Yook, Allen Ka Loon Cheung, Yong-Su Jin, Hyejin Youn, Pan-Jun Kim, Cheol-Min Ghim

**Author notes:** Correspondence and requests for materials should be addressed to H.Y., P.-J.K., or C.-M.G. These authors contributed equally (Junghun Chae, WooJoong Kim, Woochul Jung, Dawoon Jeong, Roktaek Lim).

## Abstract

Long-term innovation depends not only on discovering new elements but also on facilitating new recombination at the time of their co-availability. This notion of combinatorial innovation raises the question about whether scientific exploration and industry research persist over similar time horizons to meet innovative opportunities. Here, we analyze about 20 million papers and US, Chinese, and European patents to compare the courses of scientific and industrial activities in genetic research, a domain of far-reaching societal importance. We observe that research on new genes has declined since the early 2000s, but the exploration of novel gene combinations remains active for biotechnology innovation. Yet, fields of highly practical or commercial focus are less likely to test novel gene combinations. Furthermore, industry R&D on each gene tends to decline earlier than scientific interest in that gene. After this decline of the industrial momentum, continuous scientific research still sustains exploratory and innovative opportunities, involving novel gene combinations, clinical trials, and technologically-impactful work. Based on a hypothetical scenario of over-aligned science and industry, we estimate that up to 42.2–74.4% of the combinational opportunities would be unexplored if scientific research becomes hindered due to industry’s retreat. This study suggests the potential complementary roles of scientific and industrial research with different temporal modes in innovation cycles, calling for balanced policies to foster innovation ecosystems.

## 1. Introduction

Much of today’s technological frontier—including artificial intelligence, robotics, mRNA vaccines, and quantum computing—is advanced by industry, with a vast share of resources and research infrastructure [1–3]. This industrial activity is essential to translate discoveries into practice, yet it seems to run on a different clock from that of long-term scientific research. Specifically, each organization weighs the exploration of uncertain, distant possibilities and the exploitation of proximate, well-understood elements [4, 5]. This tension between exploration and exploitation shapes the time horizon of innovative activity: the more weight on the exploration (exploitation), the longer (shorter) this time horizon. Multiple factors underlie their balance, including the cost and return of research and development (R&D), appropriability, regulatory barriers, risk level, and the demand for timely, marketable outcomes (i.e., commercial pressure) [6–11]. For example, stronger commercial pressure likely leans toward exploitation and shorter-horizon research. Corporate R&D is frequently related to near-term opportunities and subject to investment cuts when expected returns or management conditions change [7–10]. The storied Bell Labs, an industrial laboratory once known as an innovation powerhouse across fundamental and applied sciences, could not maintain its standing amid corporate downturns and restructuring [12].

However, such industrial R&D decline, marked by reduced investment and patents, should not be read as a sign that an entire field has passed its peak of innovation. Given that innovation depends not only on discovering new elements but also on facilitating new recombination at the time of their co-availability [13–16], academia and other sectors beyond industry may still continue scientific exploration and meet the sudden opportunities ready for the combinatorial innovation to occur, until true obsolescence [17, 18]. We arrive at this possible scenario of different temporal modes of innovative activity, unlike the prior science-of-science studies that often focus on the flow of scientific or technological knowledge into future innovation [13, 19–21].

Tracing both scientific research and industry-driven technology development, we here examine the possible mismatch between their active time windows and its implications for overall innovation courses. For example, if industry’s drive begins to wane but innovative exploration would continue with longer-horizon scientific research, interpreting this time point from the industry as the true onset of obsolescence may lead to a policy mistake with the premature withdrawal of research support, a self-fulfilling prophecy. Whether scientific exploration indeed outlasts industrial momentum is thus a valuable question to empirically test, given the existing concerns that innovative research output has contracted across different domains [22, 23] and the pharmaceutical industry is risk-averse with underinvestment in novel drug development [7, 24]. A recent COVID-19 R&D study further suggests that non-monetary incentives may in fact be common drivers of innovation [6].

For our investigation, we target empirical patterns in the development of biotechnology— specifically, genetic research. Research on genes and gene products forms the foundation of modern biotechnology and is recognized for its applicability in medicine, agriculture, food industry, energy supply, environmental remediation, and grand challenges such as pandemics and climate change [24–30]. The recent year (2023) marked both the 70^th^ anniversary of the discovery of the DNA double helix and the 20^th^ anniversary of the completion of the Human Genome Project [24, 25], with an expectation for the continued exploration of breakthroughs. For simplicity in our study, both genes and gene products (proteins) will henceforth be referred to as genes, unless specified.

## 2. Results

### 2.1 Popular genes in scientific research and technology inventions

We traced research of 53,484 different genes for protein production through the systematic survey of 16 million papers and 2 million patent families, the latter from the U.S. Patent and Trademark Office (USPTO), the China National Intellectual Property Administration (CNIPA), and the European Patent Office (EPO). The genes were curated from the UniProt Knowledgebase (UniProtKB)/Swiss-Prot (Sect. 4.1) [31]. Our analysis focuses on the periods of 1990–2021 for paper publications (i.e., until the COVID-19 pandemic era) and 1990–2020 for patent applications. These periods are chosen because the Human Genome Project was launched in 1990 as a milestone in the history of genetic research, whereas the patent application records after about 2020 did not seem fully accessible at the time of this manuscript preparation (Sect. 4.2).

Like previous studies, we view papers as carriers of scientific research and patents as carriers of technology inventions [13, 19, 20, 32–35]. Accordingly, we obtained the annual counts of papers and patents for each gene as the activity levels of scientific research and technology inventions with that gene, respectively (Sect. 4.3). Because not all genes in the full text of a given document might directly contribute to its subject, we only considered genes in the title and abstract. On the other hand, our analysis shows that patents are typically company-assigned with a probability of 76.0% for each gene, whereas papers rarely include company-funded research, with only a 5.2% probability for each gene (Sect. 4.4).

The time-series of paper and patent counts thus provide the informative, albeit imperfect, profile of scientific research without direct industrial support and that of industry-driven technology inventions, respectively.

These time trajectories reveal the dynamic trends in genetic research. Hemoglobin, for example, saw its scientific research grow until 2021 with five times as many paper publications as in 1990. Similarly, hemoglobin-involving inventive activities grew with ten times as many patent applications as in 1990 (Fig. 1A). The examples of CRISPR associated protein 9 (Cas9), angiotensin-converting enzyme 2 (ACE2), and cyclooxygenase-2 (COX-2) are presented in Fig. 1B–D. Intriguingly, ACE2 is a binding target of severe acute respiratory syndrome coronavirus 2 (SARS-CoV-2) to enter host cells and cause COVID-19 [29, 30], and as expected, its research and inventive activities skyrocketed from 2020 (Fig. 1C). On the other hand, COX-2 showed the decline in the inventive activities in the 2000s (Fig. 1D), and this case will be elaborated on later. As an alternative way, among the multiple genes in a given paper or patent, we identified only the genes of the main focus in that paper or patent (Sect. 4.3). In this study, when analyzing only those main genes, we will explicitly state it.

**Fig. 1.**
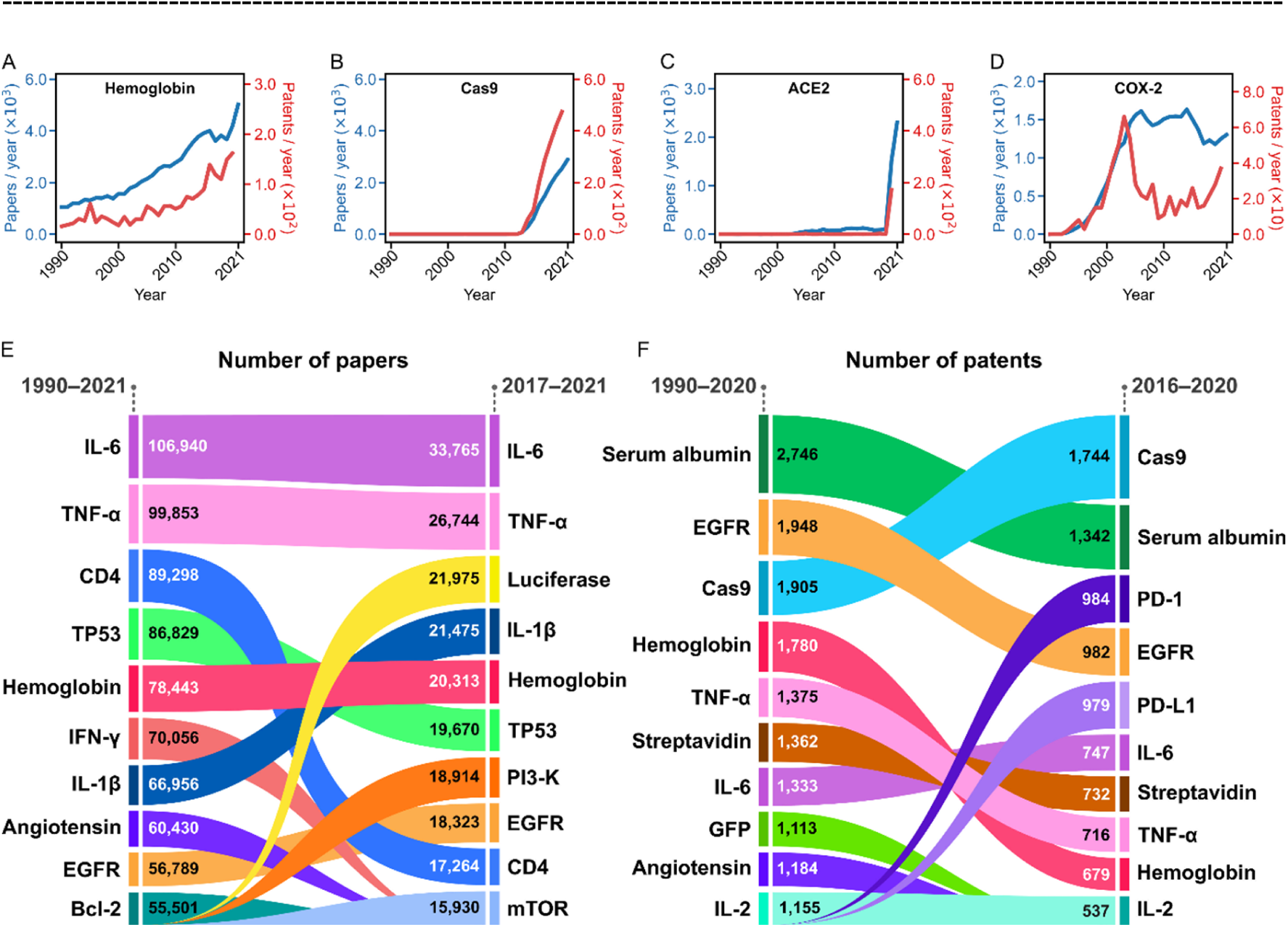
Scientific and technological trends in genetic research since the launch of the Human Genome Project (1990). **A**–**D**, Annual counts of papers (blue) or patents (red) whose titles/abstracts include a given gene. **E**, **F**, Ten most frequent genes in papers (**E**) or patents (**F**) of all time (1990– 2021 for papers and 1990–2020 for patents; left side) and latest five years (2017–2021 for papers and 2016–2020 for patents; right side). The reason for selecting different periods for paper and patent counting is explained in Sect. 4.2. Genes are vertically ordered by their paper or patent number in a given period. Each gene is indicated by a ribbon with a vertical width, proportional to the paper or patent number.

We found that interleukin-6 (IL-6) and tumor necrosis factor-alpha (TNF-α) are the top-ranked genes in the cumulative volumes of scientific research (Fig. 1E). The top ten genes for research included many immune genes, several cancer-related genes such as tumor suppressor p53 (TP53) and epidermal growth factor receptor (EGFR), and others (Fig. 1E). In the case of genes in inventive activities, serum albumin and EGFR were the top-ranked, but their positions were overtaken by Cas9 and programmed cell death protein 1 (PD-1) during 2016–2020 (Fig. 1F). PD-1 and programmed cell death ligand 1 (PD-L1) together came to the front in this period and they are both immunotherapy-related genes. Similarly, the top genes among only the genes of the main focus in papers or patents are presented in Supplementary Fig. S1 of Additional file 1 (Sect. 4.3).

These paper and patent records of genes provide an empirical basis for analyzing the scientific and industrial activities over time, as we will later unfold their global picture beyond the above prominent genes.

### 2.2 Research on new genes slows down

One may consider the pace of studying new genes as one dimension of innovation vitality in genetic research. We observe that the total number of newly researched genes in papers and patents each year, which were not homologous to any genes in prior studies, grew until 2002 and then turned to robust decline (Fig. 2A and Sect. 4.6). A similar tendency has also been observed in a previous study with only human protein-coding genes [28].

**Fig. 2.**
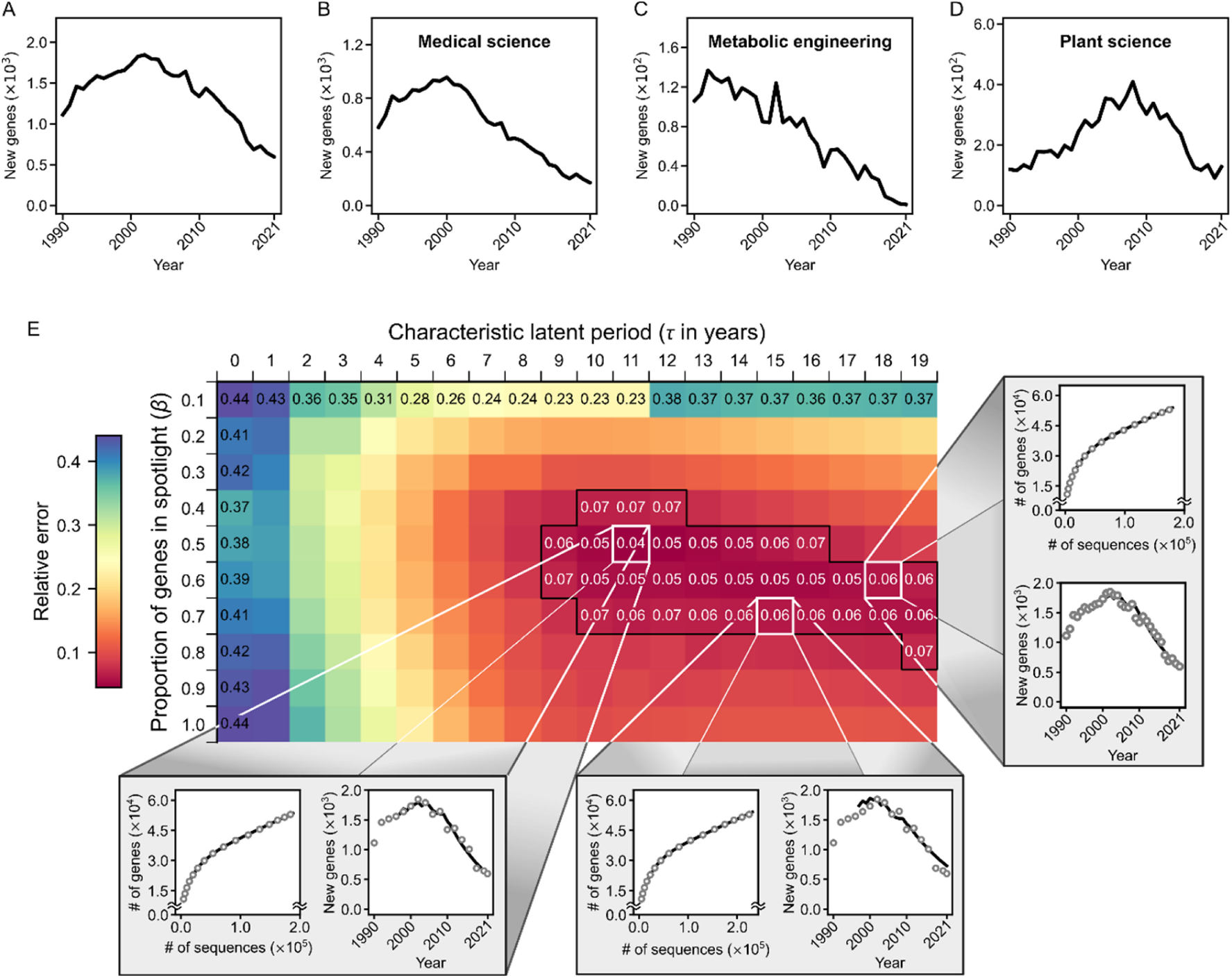
Research on new genes. **A**, Annual counts of new genes in papers and patents. **B**–**D**, Annual counts of new genes in each thematic gene category (Sect. 4.5). **E**, Output of our mathematical model for the annual counts of newly researched genes. A heatmap shows the relative errors of the annual counts of the new genes between the model output and actual data over parameters *τ* (horizontal) and *β* (vertical), while the model assumes that fraction *β* of newly reported amino acid (AA) sequences would be studied sometime later with characteristic latent period *τ* (Sect. 4.7). A parameter region with small relative errors (≤ 0.07) is enclosed by a black boundary. For particular *τ* and *β* with the small relative error, the heatmap is projected to the graph of the cumulative number of the studied genes as a function of the number of their non-redundant AA sequences, i.e., individual gene copies (left) and the graph of the annual counts of newly studied genes (right), which is relevant to **A**: the linear interpolation of the model output (solid line) and the real data points (circle) are shown together in these graphs. Over a broad range of *τ* and *β*, the model produces the quantitative pattern in **A**. For further analysis of this trend, we categorized genes according to their frequent research themes. Mainly based on the expert-examined classifications of journals and patents as the venues of genetic research, each gene was assigned to an appropriate thematic category (Sect. 4.5, and Supplementary Data S1– S3 in Additional file 2–4). As will be clear, commercially-oriented categorical genes exhibit some unique patterns amid the overall new gene decline. While newly researched genes in medical science mirror the aforementioned post-millennium declining trend, new genes in the field of metabolic engineering underwent an earlier and severer decline, even dating back to 1992 (Fig. 2B,C). Of note, metabolic engineering is based on commercial and economic motives to enhance the desirable performance of living organisms by altering their biochemical pathways [36, 37]. Likewise, its parent field, applied biotechnology shows an early decline of newly researched genes (Supplementary Fig. S4, S5 for all the gene categories and the additional information on applied biotechnology and metabolic engineering cases, respectively; see Additional file 1). We will revisit these issues later. On the other hand, those in plant science just started to decline after 2008 (Fig. 2D). Interestingly, 2008 was close to the end of the NSF-funded 2010 Project that aimed to determine the function of as many genes in model plant *Arabidopsis thaliana* as possible [38]. From 2017, the further decline in new plant-science genes stopped, mainly by the contributions of the genes related to *Arabidopsis* (48.5%) and rice (16.6%) (Fig. 2D and Sect. 4.6).

Consequently, the exploration of new genes in research and inventive activities continued to slow down soon after the turn of the millennium, which saw the announcement of the first draft human genome and the near-end of the Human Genome Project. Compared to 2002, the new genes fell into one-third in 2021 (Fig. 2A). Despite this overall tendency, an exception is the continuous increase of new genes in Chinese patents each year, unlike US and European patents (Supplementary Fig. S2A–D in Additional file 1). This exception resulted from the unusual growth of the total annual patents themselves from China. Indeed, whichever of the US, Chinese, and European patents are analyzed, the *proportion* of patents with new genes (i.e., the probability of new gene occurrence per patent) began to decline just before 2000 and finally recorded 3.3–5.4% (Supplementary Fig. S3 in Additional file 1). Such a declining trend and the other key results throughout this study remain unchanged when we only consider the genes of the main focus in papers or patents (Sect. 4.3). The overall decline in newly researched genes calls to mind the pre-existing concerns about conventionally-biased human gene studies and drug developments [7, 24].

To account for the overall decline in newly researched genes from 2002 in Fig. 2A, we developed a mathematical model that describes the cumulative number of distinct genes in papers and patents over time (Sect. 4.7). In this model, an effectively finite pool of genes and their copies across species becomes a target of current-day genetic research. For these gene copies, we considered all the possible orders of their initial studies with approximately equal probabilities, and then calculated the expected cumulative number of the distinct studied genes each year in combination with annually-updated gene copy numbers from the UniProtKB/Swiss-Prot data. This model accurately reproduces Fig. 2A and also gives other results compatible with empirical data over a broad range of conditions (Fig. 2E, and Supplementary Fig. S6 in Additional file 1; *P* <10^−4^ and Sect. 4.7). For any given type of model output, we accepted the collection of all the individual outputs from those feasible model conditions, not the particular ones. Our model suggests that the vast majority (71.3 ± 5.4%) of taxonomically-dispersed genes studied as of 2021 had already been studied at least once before 2002 (avg. ± s.d. and Sect. 4.7). In the case of taxonomically-restricted genes [39, 40] as of 2021, only their 8.0 ± 0.8% had been studied before 2002, though (avg. ± s.d.). Hence, the time around the completion of the Human Genome Project was a turning point with the substantial accumulation of the researched, taxonomically-dispersed genes, which then rarified the exploration of the new ones. Yet, we do not exclude the possibility that the decline in the studies of new genes in Fig. 2A may later be stopped by rapidly-growing breakthrough technologies such as third-generation sequencing, although this sequencing still remains minor in its adoption (Sect. 4.8, and Supplementary Fig. S7 in Additional file 1).

### 2.3 Novel gene combinations keep actively explored, but not in every field

Despite the significance of new genes, a more common mode of novel genetic research could be the use of known genes for new combinations. In stark contrast to the slowdown of studying new genes from 2002, new combinations of genes tended to increase until recently (Fig. 3A and Sect. 4.9). There exist some variations between different patent sources: from the 2000s, the numbers of novel gene combinations in Chinese and US patents were accelerating and running steadily, respectively. Still, none of US, Chinese, and European patents entered a decline phase in novel gene combinations and they all remained active for those combinations (Supplementary Fig. S2E–P in Additional file 1).

**Fig. 3.**
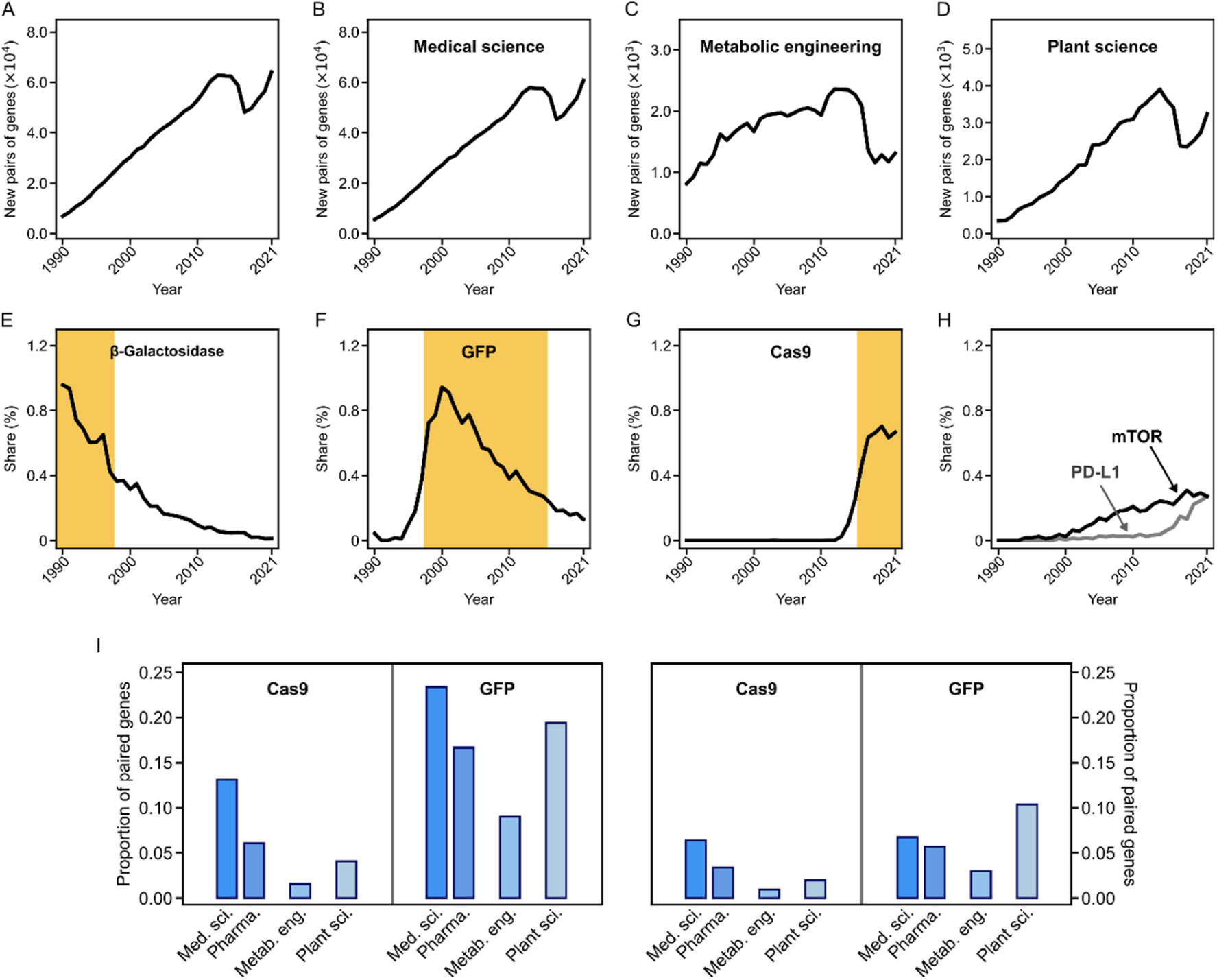
Research on novel gene combinations. **A**, Annual counts of new pairs of genes in papers and patents. For separate counting of paper and regional patent cases, and for larger gene combinations beyond a pair, see Supplementary Fig. S2E–P in Additional file 1. **B–D**, Annual counts of new pairs of genes in each thematic gene category (Sect. 4.9). **E–G**, For each gene that once had the highest share (relative frequency) in new pairs of genes, its share is presented over time. Shaded is the period when this gene had the highest shares. **H**, The shares of PD-L1 and mTOR in new pairs of genes over time. They were not the highest but still ranked high. **I**, The proportion of genes in each thematic gene category, paired with Cas9 or GFP until the peak time of its annual paper and patent counts. In this pairing, we considered genes that debuted in papers/patents in 1990–2000 (left) and 2001–2011 (right), separately (Sect. 4.9). Abbreviations: med. sci., medical science; pharma., pharmaceutical; metab. eng., metabolic engineering; plant sci., plant science.

Regarding different thematic categories of genes, we observe an increase in novel gene combinations in medical science and a similar but recently weaker trend in plant science. About commercially-oriented categories, we observe a contrast in metabolic engineering and applied biotechnology with drop-offs and stagnancy in the novel gene combinations from 2017, possibly warning for their low innovation vitality (Fig. 3B–D, and Supplementary Fig. S8 in Additional file 1 for all the gene categories; see also Sect. 4.9).

Back to the overall trend beyond regional and categorical patterns, this “innovation by combination” [13–16] involves the succession of highly frequent genes in those combinations over time: the most frequent in the newly paired genes were β-galactosidase in 1990–1997, green fluorescent protein (GFP) in 1998–2015, and Cas9 in 2016–2021, along with less yet highly frequent genes such as mammalian target of rapamycin (mTOR) and PD-L1 (Fig. 3E– H, and Supplementary Table S1 in Additional file 1). For example, Cas9 newly teamed with 4,141 different genes between 2016 and 2021. We notice that the genes of such frequent fresh partnership have been the iconic workhorses of biotechnology innovation, recapitulating the historical trajectory of landmark ideas and technologies from DNA cloning (β-galactosidase) and live cell imaging (GFP) to genome editing (Cas9) (Fig. 3E–G) [41, 42].

Among the above prominent genes of biotechnology innovation, Cas9 and GFP are recognized for their utility across disparate domains such as medical and plant sciences. Therefore, we examined how many genes in each category teamed with Cas9 or GFP across papers and patents, especially during the early stages of Cas9 and GFP. We found that, in their infancy to peaks, Cas9 and GFP respectively teamed with 6.4–13.1% and 6.7– 23.4% of medical science genes that debuted in papers or patents during similar periods (Fig. 3I and Sect. 4.9). Interestingly, if we limit our focus to pharmaceutical genes, they fall short of this gross medical science result, with the particularly lower proportion for Cas9 (Fig. 3I; *P* ≤ 2 ×10^−4^ and Sect. 4.9). Of note, pharmaceutical research within medical science is known to prioritize marketable outcomes. Another observation from our data is that both metabolic engineering and (its parent) applied biotechnology rank among the lowest in the proportions of the genes teaming early with Cas9 or GFP (Fig. 3I, and Supplementary Fig. S9 in Additional file 1 for all the gene categories; *P* ≤10^−4^ and Sect. 4.9). These patterns do not much depend on specific periods of the early teaming (Supplementary Fig. S9 in Additional file 1). The patterns of pharmaceutical, metabolic engineering, and applied biotechnology fields indicate that the highly practical focus of these fields may be in fact limiting the test and adoption of newcomer technologies. There exist further field-specific issues, such as the emphasis on the price competitiveness of products in the cases of metabolic engineering and applied biotechnology, discouraging high-risk exploratory approaches. Such exploration restraint appears to be consistent with the previous, long-alarming trends in Fig. 2C, 3C and Supplementary Fig. S4D,E, S8D,E (Additional file 1), which exhibited the severe decline of both newly researched genes and novel gene combinations in metabolic engineering and applied biotechnology. Next, we will further elaborate on the time horizon of commercially-oriented, industry-relevant research activities.

### 2.4 Short-horizon industrial drive and long-horizon scientific activity for innovation

Recall the very initial finding in this study that patent-filed technology inventions are typically industry-driven, while paper-published scientific research is not. Industry R&D is thus mainly imprinted on patent trajectories, and our subsequent results do remain valid when considering only the company-assigned patents among all the patents.

Fig. 4A shows the paper and patent time-series of COX-2, which has been studied for anti-inflammatory treatment and pain relief [43–47]. Notably, its annual patent applications started to decline in 2004 while the paper publications still persisted (Fig. 4A). This patent falloff, in fact, coincided with the withdrawal of a COX-2 inhibitor drug for arthritis patients (Vioxx) from the market, due to the risk of heart attack and stroke [43]. Phosphodiesterase 5 (PDE5) from our data, well known for its inhibitor sildenafil to treat erectile dysfunction (Viagra), is another gene with a patent falloff but persistent paper publications. The use of sildenafil was approved for pulmonary arterial hypertension in 2005, following its previous approval for erectile dysfunction [48, 49]. Upon sildenafil’s dominance in the market, patent applications with PDE5 started to fall in 2007. Therefore, the cases of COX-2 and PDE5 presumably exemplify the industry’s R&D depression due to risk management and anticipated lower returns in a saturated market, respectively [7, 9, 10].

**Fig. 4.**
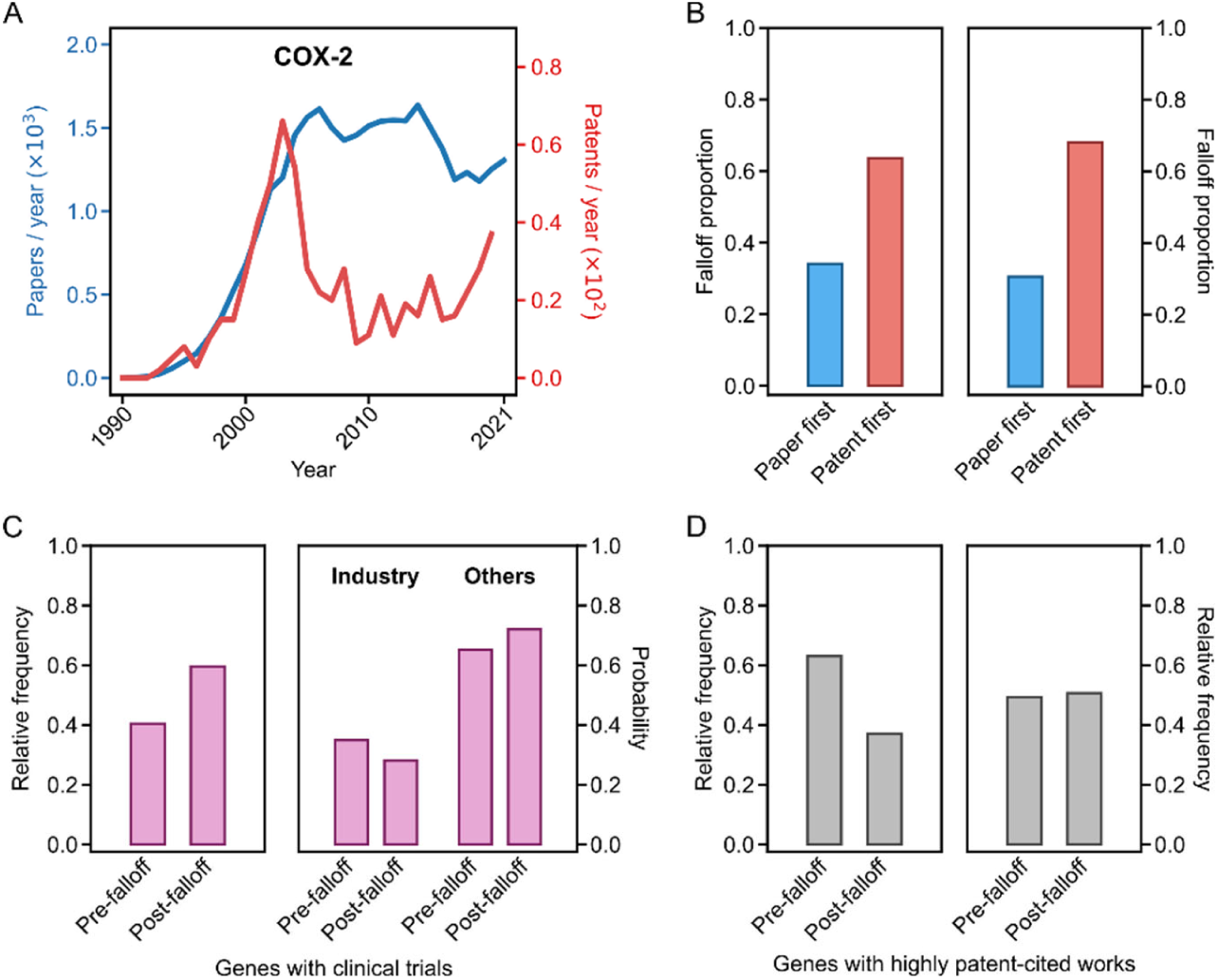
Paper–patent falloff relationship and research continuation. **A**, Annual paper (blue) or patent (red) counts of COX-2. **B**, The proportion of genes whose paper falloffs precede the patent falloffs (blue) and the proportion of genes whose patent falloffs precede the paper falloffs (red). Here, we considered genes that debuted in papers/patents in 1990–2000 (left) and 2001–2011 (right), separately. For a fair comparison between paper and patent falloffs, we set the endpoint of the paper time-series to the same year (2020) as the patent time-series. **C**, On the left side, the proportion of genes with clinical-trial papers before (“Pre-falloff”) or after/at (“Post-falloff”) their patent falloffs, among the clinical-trial genes that debuted in papers/patents in 2002–2011. These debut years were chosen regarding the first year of the mandated trial posting to ClinicalTrials.gov (Sect. 4.13). On the right side, the probability of industry or other sponsorship for a clinical trial with each gene in a given case (“Pre-falloff” or “Post-falloff”) from the left side. **D**, The proportion of genes with technologically-impactful papers before (“Pre-falloff”) or after/at (“Post-falloff”) their patent falloffs. These papers received ≥3 patent citations each within the first three years of publications. We considered the genes that debuted in papers/patents in 1990–2000 (left) and 2001–2011 (right), separately. Alternative criteria for high patent citations did not qualitatively change the observed result here.

From now on, we define the paper or patent’s falloff as a robust decrease of its number by at least a quarter of the peak value. If the paper or patent volume of a given gene was too small for this definition, we inferred the falloff from a dense presence-to-scarcity transition (Sect. 4.10). Alternations of the falloff definition did not change our main results.

Remarkably, the preceding patent falloff above is not limited to COX-2 or PDE5 but characterizes the general course of genetic research. For two-thirds of genes that debuted in papers or patents during similar periods, their patent falloffs preceded the paper falloffs (Fig. 4B; *P* < 10^−4^ and Sect. 4.10). For example, genes that debuted in 1990–2000 show their patent falloffs 6.5 ± 4.9 years (avg. ± s.d.) ahead of the paper falloffs, and a quarter of these genes even show ≥10-years earlier patent falloffs. Our further analysis excludes the possibility that the policy requirement of non-overlap of a patent content with prior art [11] might contribute to the rapider patent falloff (Supplementary Fig. S10 in Additional file 1, and Sect. 4.10). We hence confirm that industry R&D on a given gene tends to have a shorter time horizon than the scientific interest in that gene. One natural consequence of this relatively early patent falloff is that the total patent volume of each gene becomes capped by the total paper volume with a certain scaling (Supplementary Fig. S11 in Additional file 1, and Sect. 4.11).

During the period of continuous, long-horizon scientific research without much of the industry’s drive, the pursuit of breakthroughs and impactful works did not stop as we will now demonstrate. Fig. 4C shows that even ∼60% of the genes associated with clinical trials underwent these trials during the times of their limited patent applications (Sect. 4.13).

Besides, trials with each gene were usually not sponsored by the industry, with a ∼70% probability whether conducted during the limited patent applications or not (Fig. 4C; *P* ≤ 0.05 and Sect. 4.13). This fact reminds us of the previous report that 70% of COVID-19 clinical trials were driven by public research institutions [6]. To evaluate the technological impact of scientific research, we measured it by the patent citations of each paper within a defined period [13, 16, 19, 20, 32, 34]. We found that a substantial body of the genes (37.0–50.7%) with the highly impactful papers did appear in those papers during the times of the limited patent applications (Fig. 4D and Sect. 4.14).

These general tendencies are repeated in the cases of COX-2 and PDE5. After the patent falloff, COX-2 continued to be studied for its function in various research papers, along with many clinical trials [44–47, 50]. They include the studies of the COX-2, EGFR, and mTOR relations in cancer, later followed by the clinical trial [44, 45], the relationship between ω−3 fatty acids and COX-2, with many patent citations [46], and the relief of hyperinflammatory responses in COVID-19 [47]. In the case of PDE5 after the patent falloff, the PDE5 inhibition continued to be tested in paper-published research and clinical trials from various aspects, such as β-catenin-involving anti-cancer mechanism and muscle ischemia alleviation, both followed by the patent citations [51–53], cardiovascular risk reduction [54], and ventilation improvement for single-ventricle patients [55]. In other words, the industry’s interest in these genes turned cold first, but the innovative scientific research persevered with longer time horizons.

### 2.5 Long-horizon scientific research maintains exploratory opportunities

Our results indicate that genes with the fading industry’s interest still retain the opportunities to explore innovative approaches and possible breakthroughs. To further examine this issue, we preselected eight iconic genes in biotechnology innovation, based on their shares in novel gene combinations explained in Sect. 2.3: Cas9 for genome editing [42], PD-L1 for immunomodulation and immunotherapy [56], mTOR and TP53 mainly for cancer research [57, 58], GFP and luciferase for live cell imaging [59, 60], β-catenin mainly for regenerative medicine and cancer research [61, 62], and β-galactosidase for DNA cloning [41] (Sect. 4.12). Hereafter, we will just refer to these genes as *iconic genes*. Interestingly, the aforementioned COX-2 teamed with two iconic genes—PD-L1 and Cas9—for the first time after its patent falloff, although the patent applications reduced to one-third and one-fourth of the peak, respectively (Fig. 5A). In these cases, PD-L1 and Cas9 had not yet reached their own peaks, but they respectively served for the investigations of the suppression of transplant rejection and the mechanism of gastric cancer [63, 64]. Likewise, the aforementioned PDE5 newly teamed with three early-stage iconic genes—GFP, TP53, and mTOR—when its patent applications reduced to one-third to one-fourth of the peak. GFP, TP53, and mTOR here served for the monitoring of cyclic guanosine monophosphate (cGMP) levels inside living cells and the investigations of kidney injury protection and liver cancer treatment, respectively [65–67].

**Fig. 5.**
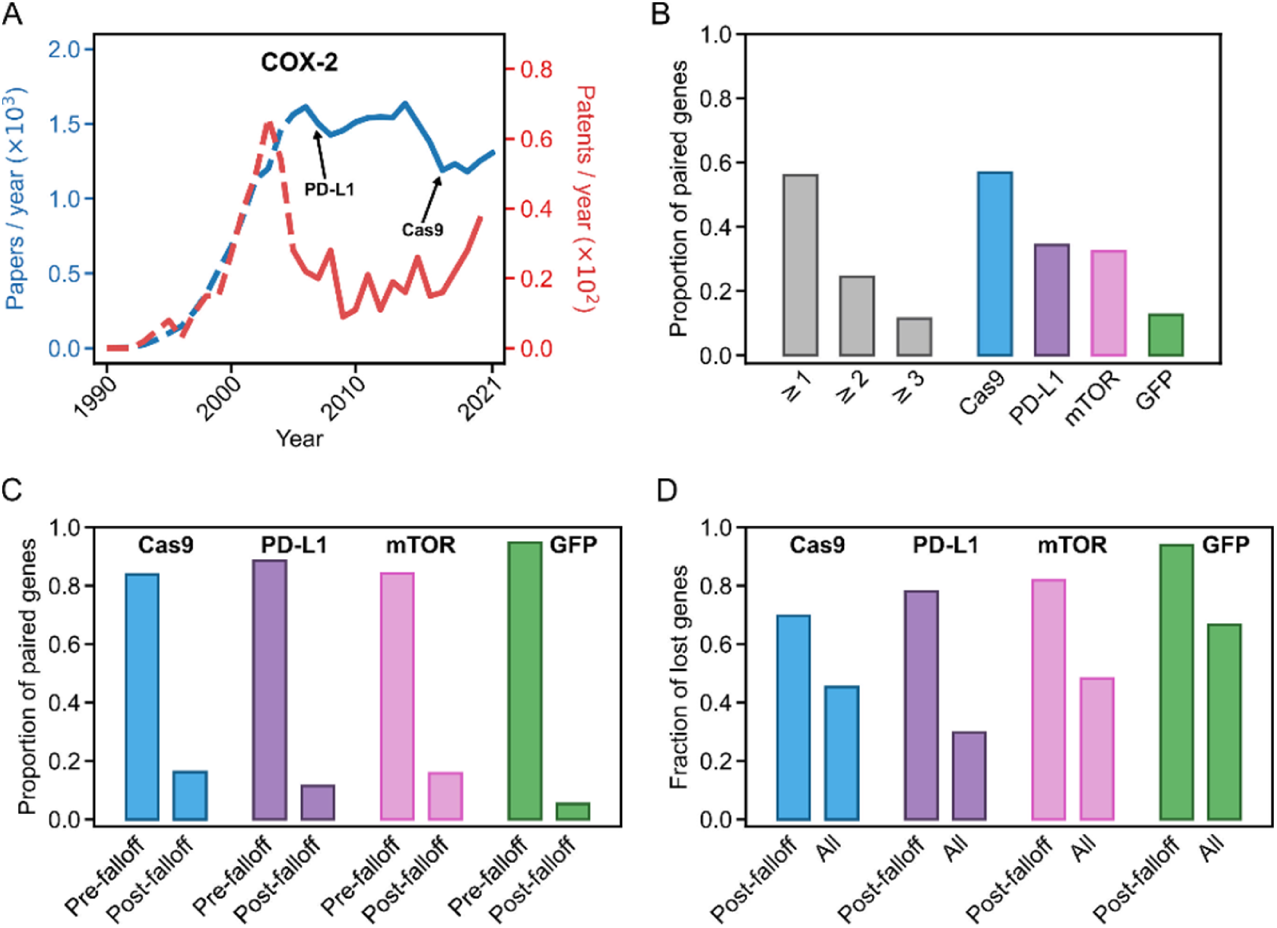
Teaming with new iconic genes in their early stages. **A**, Annual paper (blue) or patent (red) counts of COX-2 (dashed line for a period before its patent falloff). An arrow indicates the initial time of the teaming with each iconic gene if this teaming comes after or coincides with the patent falloff. **B**, The proportion of genes teaming with ≥1, ≥2, or ≥3 new iconic genes after (or at) the patent falloffs, and the proportion of genes initially teaming with a given iconic gene—Cas9, PD-L1, mTOR, or GFP—after (or at) their patent falloffs (Sect. 4.15). We only considered the teaming with the iconic genes until the peak times of their annual paper and patent counts. **C**, The proportion of genes initially teaming with a given iconic gene before (“Pre-falloff”) or after/at (“Post-falloff”) their paper falloffs, among the genes with this teaming after/at the patent falloffs (Sect. 4.15). We only considered the teaming with the iconic gene until the peak time of its annual paper and patent counts. **D**, The estimated proportion of genes that would lose the teaming with a given iconic gene, among the genes that initially teamed with this iconic gene amid their limited patent applications (“Post-falloff”) or among all the genes that teamed with this iconic gene (“All”) (Sect. 4.16). We only considered the teaming with the iconic gene until the peak time of its annual paper and patent counts. In **B**–**D**, we considered genes that debuted in papers/patents in 1990–2000. Genes with post-2000 debuts were not considered due to the fundamentally low overlap of their research periods with the early stages of the iconic genes.

Indeed, we found that a substantial portion of genes teamed with new iconic genes after their patent falloffs, although these iconic genes were still in the early stages of work. Of genes that teamed with iconic genes in their infancy to peaks, the majority (54.5%) initially teamed with at least one of these iconic genes after the patent falloffs (Fig. 5B and Sect. 4.15). About a half of these genes, including COX-2 and PDE5, initially teamed with two or even more of those iconic genes after the patent falloffs (Fig. 5B and Sect. 4.15). In addition, for one-third to a half of the genes that teamed with Cas9, PD-L1, or mTOR in its infancy to peak, these teams were initially formed after the patent falloffs, and this fraction had already been appreciable during an earlier period of the teaming (Fig. 5B, Supplementary Fig. S12 in Additional file 1, and Sect. 4.15).

These results suggest that the loss of the industrial momentum does not necessarily close the exploratory opportunities of genetic research. Further analysis reveals that sustained, long-horizon scientific research is a key to bring this possibility to life, despite the fading industry’s interest. Specifically, the teaming of each gene with a new, early-stage iconic gene after the patent falloff predominantly occurred (≥72.9%) when the active paper publications still continued (Fig. 5C, and Supplementary Fig. S13 in Additional file 1; *P* <10^−4^ and Sect. 4.15).

These findings raise a fundamental question: if the turnover of industrial targets propagates to scientific activities outside the industry through financial or other impacts [9, 68], to what extent would it disturb the exploratory opportunities of genetic research? One instance of such propagation is the blow to biofuel research at leading universities in the mid-2010s due to the oil price drop and subsequent investment cuts [69–71]. To answer our question, we conducted a “thought experiment” with a hypothetical scenario that scientific research on a given gene becomes hindered by a lack of the industrial momentum: specifically, we altered the temporal trend of the paper counts to match that of the patent counts, which had come to a low level (Sect. 4.16). The lack of the industry’s visible interest in a particular research area may not only affect its own investment but also possibly diminish the public and governmental support for that research area [72]. Our thought experiment gives rise to a remarkable estimation that the vast majority (≥ 69.7%) of the genes, which initially teamed with a given early iconic gene amid the limited industrial momentum and patent applications, would now lose this teaming (Fig. 5D, Supplementary Fig. S14 in Additional file 1, and Sect. 4.16). Among all the genes whether affected or not by our hypothetical scenario, still 42.2–74.4% of the genes that once teamed with a given early iconic gene would lose this teaming, with the exception of PD-L1 although it would also lose one-fourth of the teams (Fig. 5D, Supplementary Fig. S14 in Additional file 1, and Sect. 4.16). In the examples of COX-2 and PDE5 whose annual patents fell in 2005 and 2007, the hindering of their scientific research would drastically reduce the chances of the teaming with each early iconic gene by ≥ 83.5% at most (Sect. 4.16). Hence, these estimations highlight the cost of over-aligning with industry-driven research, which would miss the long-term exploratory and innovative opportunities once the industrial momentum wanes.

## 3. Discussion

Through a decades-long overview of genetic research, our study reveals how innovative capacity is sustained despite the recurrent turnover of industry’s interest. As observed in this study, research on new genes has declined since the early 2000s, but the exploration of novel gene combinations has expanded and remains active for biotechnology innovation (Fig. 2A, 3A). This form of combinatorial innovation [13–16], however, does not evenly appear across different fields. The fields of highly practical or commercial focus, such as metabolic engineering with its industrial tie, are less likely to embrace innovative technologies with novel gene combinations (Fig. 3C,I, and Supplementary Fig. S8, S9 in Additional file 1). Consistently, the pursuit of proximate outcomes is known to favor exploitation over exploration [4, 5], and this tendency may shorten the time horizon of industry R&D. Indeed, our analysis confirms that industry R&D on each gene typically declines earlier than scientific interest in that gene (Fig. 4B, and Supplementary Fig. S10 in Additional file 1). Still, scientific research over longer time horizons tends to sustain the opportunities for exploration and innovation, involving novel combinations of genes at the early stages of their co-availability, clinical trials with genes, and technologically-impactful research (Fig. 4C,D, 5C, and Supplementary Fig. S13 in Additional file 1). We do not interpret these results as a mere dichotomy between scientific and industrial research, but as their potential complementary roles with different temporal modes in innovation cycles: industry activity helps translate discoveries into practice and raise public attention, while continuous scientific research maintains the exploration that may require more time to bring all the elements for recombination together but then possibly attract the industry again through its practical value [73–77].

Our findings indicate that an innovation strategy overly based on commercial dynamism or industrial interests will quietly endanger the exploratory research on which future breakthroughs depend. Our thought experiment does reveal the scale of this dependence on long-term scientific research: approximately, up to 42.2–74.4% of the opportunities for notable gene combinations would be unexplored if the scientific research loses its support in step with the industry’s limited interest (Fig. 5D, and Supplementary Fig. S14 in Additional file 1). Therefore, a proper role of public funding would be stable support for perseverant, exploratory science [74–76] even in the areas that have already passed the peaks of the industrial drives. Meanwhile, the government may also encourage the industry to pursue exploratory, longer-horizon innovation alongside near-term commercialization [12, 78]. One way could be the governmental partnership to de-risk the commercialization of new technologies by offering the opportunities to test, apply, and publicize them [79, 80] and helping establish the communication channels with consumers to predict market response [81, 82].

Although our data analysis takes various controls and null models to minimize the possibility of alternative data interpretations (Sect. 4.7, 4.9–4.11, 4.13–4.16), future work should identify more specific causal relationships and mechanisms. This identification can benefit from the use of highly-refined paper and patent data [83–85] with affiliation, citation, funding, investment, licensing, and firm-level R&D records. Consideration of hidden biases in patent data due to commercial interests is also warranted [21]. Our analysis could be further extended to additional forms of innovative genetic research, such as the discovery and studies of previously-known genes within unexpected taxa [86].

In conclusion, new opportunities will come when explored long enough for notable combinations and innovations to take shape. Industrial momentum advances the opportunities toward real life over short time horizons, and scientific research keeps them open even after the industry’s retreat. Recognizing this temporal division of exploratory labor is the basis of constructive policy—not to maximize any single mode of research, but to foster an innovation ecosystem where short- and long-horizon activities reinforce each other. This view will be essential to unlock the full potential of biotechnology and to address complex challenges that require persistent experimentation across uncertain and evolving knowledge landscapes [1, 87–89].

## 4. Methods

### 4.1 Listing of gene names

We used UniProtKB/Swiss-Prot as the source of gene and protein names. Each entry of the UniProtKB/Swiss-Prot database presents the information of an expertly curated, non-redundant protein sequence mainly at the species level, derived from predicted protein domains, systematic rule-based annotation, and careful proteome clustering [31, 90]. For example, COX-2 protein has the entry for each species such as human (*Homo sapiens*), mouse (*Mus musculus*), rat (*Rattus norvegicus*), or horse (*Equus caballus*). One entry for the COX-2 protein includes the names/synonyms such as cyclooxygenase-2, prostaglandin G/H synthase 2, prostaglandin H2 synthase 2, and PTGS2. Across all the UniProtKB/Swiss-Prot entries, we collected gene and protein names and their synonyms from the attributes in Supplementary Table S2 of Additional file 1 (UniProtKB/Swiss-Prot XML file, Release 2022_04). These names and synonyms were later used for their search in paper and patent texts. For simplicity, we here refer to a set of homologous genes and their proteins as a gene, and refer to their names and synonyms as synonyms unless specified. The comprehensive synonym set of each gene was identified by the grouping of the entries with highly shared synonyms, as these entries often corresponded to homologous genes.

Specifically, we excluded improper synonyms such as the names of cleaved chains or subdomains of main peptide bodies, and constructed the graph where each entry is a node and two nodes are linked if their entries share a substantial portion of the synonyms to satisfy |*A*∩ *B*|⁄min(|*A*|, |*B*|) ≥ 0.75 (*A* and *B* are the synonym sets of these entries). By applying the generalized Louvain method to this graph (v0.3; https://sourceforge.net/projects/louvain) [91], we detected densely-knitted subgraphs, i.e., network communities. For synonyms still shared by multiple network communities, each synonym was assigned only to the community with the largest share of the entries for that synonym. The whole synonyms in each community were then considered the gene’s tentative synonym set. After the test run of synonym matches with paper and patent texts (Sect. 4.3), we identified some homonyms irrelevant to genes. To reduce these homonyms, we removed non-biological phrases in common use based on their frequent occurrences in published books in society, available from *Google Books Ngram Corpus* (version 3, February 2020) [92]: the phrases with higher counts than a heuristically-chosen threshold (1.8×10^7^ counts in the books published by 2020) were prioritized for removal from our gene synonyms. These prioritized ones and the remaining synonyms, particularly around the threshold, underwent extensive manual inspection to reliably exclude non-biological phrases and retain the relevant terms. We then curated and finalized our gene synonym sets.

### 4.2 Collection of paper and patent data

We created complementary, paper and patent corpora to survey research and inventive activities in genetic subjects, respectively.

In the case of papers, we targeted those in the journals of biology, applied biology, medicine, and other related disciplines. The categories of those disciplines and the associated journal lists were obtained from the Web of Science (WoS) Journal Citation Reports (JCR) (https://jcr.clarivate.com). We manually curated these journal lists for better association with the journal categories and removed the journals irrelevant to contemporary genetic research. The resulting journal lists are provided in Supplementary Data S1 with our expert-curated journal categories JG0, JG1, ···, and JG8 (Additional file 2): JG1 includes biomedical journals, along with its subsets JG2 and JG3 for veterinary medicine and pharmaceutics, respectively. JG4 includes journals in applied biotechnology, along with its subset JG5 for metabolic engineering. JG6 and JG7 include plant science and general microbiology journals, respectively (we here use the term “general microbiology” to distinguish it from applied microbiology covered by JG4). JG0 includes all the journals in JG1 to JG7, with additional journals for animal, algal, and aquatic research. JG8 includes broad-scope journals and other journals not fit to JG0.

At the same time, we downloaded the annual releases of the baseline data files from the PubMed FTP server (April 22, 2022; https://ftp.ncbi.nlm.nih.gov/pubmed/baseline) [93], which contained the titles and abstracts of the articles serviced by PubMed. Other than the titles and abstracts, we did not pursue the full text analysis because all the genes in the full texts were not necessarily relevant to the very subjects of the articles. We considered only the articles in the above JG0 and JG8 journals and published in 1990–2021 because the Human Genome Project was launched in 1990 and our gene synonyms derived from the UniProtKB/Swiss-Prot data dated May 2022 as noted in Sect. 4.1.

In the case of patents, we collected the USPTO, CNIPA, and EPO patent application data (US, Chinese, and European, respectively) in the PATSTAT Global database (version 5.22, 2023 Autumn Edition) provided by the EPO. For each patent family or any set of patents with duplicated titles and abstracts, we selected only the earliest-dated patent to avoid data redundancy. Among the Cooperative Patent Classification (CPC) codes jointly managed by the EPO and the USPTO (dated May 2, 2022) and the International Patent Classification (IPC) codes managed by the World Intellectual Property Organization (WIPO) (dated October 1, 2023), we selected possibly gene-related codes based on expert opinions (Supplementary Data S2 in Additional file 3) and considered the patents with these codes. There is a known delay between patent application and publication, and we did observe a roughly two-year delay with a spurious decrease in recent patent application records.

Therefore, taking this delay and our data access in Fall 2023 into consideration, the patents filed by the end of 2020 were included in our further analysis. Throughout this study, we considered only the patent titles and abstracts because the genes there mostly overlapped with those in the claims and the rest included genes not necessarily relevant to the very subjects of the patents.

### 4.3 Search of genes in papers and patents

To find the synonyms of each gene in the paper or patent titles and abstracts, we used the string-matching algorithm that directly compares the phrases in the target literature to the strings from the gene synonyms, while incorporating the indexed tree and tokenizing scheme for efficient computation [94]. Several rules were added to the search algorithms to consider common variations in the synonyms, such as the use of Greek letters, plural forms, and different letter cases. For each gene, papers and patents were counted when matching at least one of its synonyms. However, papers and patents with too many genes (>10 genes each) were excluded from our analysis, to avoid ambiguous cases with no clearly-identifiable research targets among the genes.

To systematically evaluate our gene search algorithm, we evenly sampled 100 papers and patents (with various query genes) for each test group of a gene attribute—the initial year of research in papers/patents (debut year), the total paper and patent counts, the patenting region (at least once), or the thematic gene category (Sect. 4.5). These test groups are specified in Supplementary Table S3 in Additional file 1. The precision and recall of the gene search algorithm were 0.86–0.93 and 0.66–0.76 across the test groups, respectively, based on the comparison with manually identified genes from the sampled papers and patents (Supplementary Table S3 in Additional file 1, and Supplementary Data S4 in Additional file 5). Although the existing resource, PubTator3 [95] also provides a list of genes in each article with slightly better precision (0.91–0.98), its recall is overly variable (0.18– 0.69; Supplementary Table S3 in Additional file 1, and Supplementary Data S4 in Additional file 5). Critically, PubTator3 covers only PubMed papers, and thus we excluded its use for consistency of our paper and patent analyses.

Throughout this study, we performed additional analyses that involve only the genes of the main focus in papers or patents. We identified these genes with the following heuristic rules: in the case of a paper, we considered a gene as of its main focus if this gene appears in the title or appears ≥3 times in the abstract. In the case of a patent, we considered a gene as of its main focus if this gene appears in the title or appears ≥2 times in the abstract, or if this gene is the only gene in the title and abstract. We validated these rules by the manual interpretation of 100 randomly chosen papers and patents. This manual inspection showed the precision of 0.86 and the recall of 0.90 for papers and the precision of 0.88 and the recall of 0.85 for patents (Supplementary Data S5 in Additional file 6).

### 4.4 Paper funding source and patent assignee analyses

We manually identified the funding sources of 300 different papers in genetic research, selected evenly as will be explained later (Supplementary Data S6 in Additional file 7). If the funding sources of a given paper included at least one company, the paper was considered company-funded. In the case of other funding sources from this paper set, we confirmed their public nature by reviewing the specific funding sources. For the selection of these 300 papers, we first randomly selected 50 different genes each (total 200), whether they debuted in papers/patents in 1990–2000 or 2001–2011, and whether their patent time-series include falloff years or are fully spanned by states II and III, which will be defined in Sect. 4.10.

These four cases were considered due to their high relevance to many analyses in this study. In the cases of the patent falloffs, one paper before the patent falloff and the other paper after (or at) this falloff were randomly selected for each gene; otherwise, any single paper was randomly selected for each gene. The probability of papers being company-funded for each gene was obtained by the weighted average of the proportion of company-funded papers over all those six cases, reflecting two from each patent-falloff case (one before the falloff and the other afterward). In this calculation, we assigned a weight to each case using the actual proportions of genes and their papers from the entire dataset.

For patent assignees of unambiguous characters, we considered “COMPANY”, “GOV”, “NON-PROFIT”, “UNIVERSITY”, and “HOSPITAL” among the patent applicant sectors from the PATSTAT database. For the genes with patents assigned to these sectors, we calculated the average proportion of patents, each with at least one company assignee. This quantity represents the probability of patents being company-assigned for each gene.

### 4.5 Thematic categorization of genes

The common categorization of genes based on molecular or biochemical contexts rarely allowed the best interpretation of our paper or patent counts with the genes. Therefore, we categorized the genes according to their frequent research themes. Mainly based on the journal categories and CPC/IPC codes of the papers and patents with each gene, the gene was assigned to one of the following thematic categories (Supplementary Data S3 in Additional file 4). The methods described here were heuristically developed by the examination of the categorization results.

*“Medical science”*: for each gene, we calculated the ratio of its papers in the JG1 journals in Sect. 4.2 to those in the JG0 journals (Supplementary Data S1 in Additional file 2). Genes with these ratios greater than or equal to an empirically-chosen threshold (0.7) were assigned to category “medical science”. In addition, for each gene, we calculated the fraction of its patents with CPC or IPC codes “A61K38”, “A61K31”, “A61K39”, “A61K48”, and their child codes (Supplementary Data S2 in Additional file 3). Genes with these fractions greater than or equal to an empirically-chosen threshold (0.5) were also assigned to the category “medical science”.

*“Genetic tools”*: the genes utilized as the tools for a range of molecular biology and genetic experiments were assigned to category “genetic tools” based on manual literature search and expert opinions, in the absence of their specific journals and patent classification codes. This category itself was not used for our analysis, but for excluding its genes from other categories below.

*“Applied biotechnology”*: this category was intended to include the genes mainly in non-medical applied biotechnology. For each gene, we calculated the fraction of its patents with CPC or IPC code “A61K” and its child codes (Supplementary Data S2 in Additional file 3). For each of the genes with these fractions lower than an empirically-chosen threshold (0.5), we counted its papers in the JG4 journals (Supplementary Data S1 in Additional file 2). Each gene with this count ≥ 40% of the total number of its papers in the JG0 journals was assigned to category “applied biotechnology”. To avoid the overlap with the other categories, the genes in the previous categories “medical science” and “genetic tools” were here excluded from the outset.

*“Plant science”* and *“general microbiology”*: as noted in Sect. 4.2, we use the term “general microbiology” to distinguish it from applied microbiology covered by “applied biotechnology”. For each gene, we calculated the ratio of its papers in the JG6 (JG7) journals to those in the JG0 journals, and the genes with these ratios ≥ 0.5 were assigned to category “plant science” (“general microbiology”). To avoid the overlap with the other categories, the genes in “medical science”, “genetic tools”, and “applied biotechnology” were here excluded from the outset. The overlap between “plant science” and “general microbiology” was avoided by manual recategorization of their shared genes.

A part of genes in “medical science” were subcategorized as follows: for each gene in “medical science”, we calculated the ratio of its papers in the JG3 journals to those in the JG1 journals. The genes with these ratios ≥ 0.25 were assigned to subcategory *“pharmaceutical”*. Another subcategory *“immune”*, given its definite biological function and existing database, was assigned the genes of Gene Ontology (GO) term “immune response” (GO:0006955) and its child terms, but “hemocyte proliferation” (GO:0035172) or its child terms [96]. Additionally, the genes associated with Anatomical Therapeutic Chemical (ATC) codes “immunostimulants” (L03) and “immunosuppressants” (L04) in the DrugBank database [97] were manually reviewed and thereby assigned to the subcategory “immune”. Note that the above subcategory “pharmaceutical” allows the overlap with this subcategory “immune”.

Each gene in “applied biotechnology” was assigned to subcategory *“metabolic engineering”*, when the ratio of its papers in the JG5 journals to those in the JG4 journals was ≥ 0.4.

The genes in the above categories and subcategories, particularly those near their thresholds, underwent further manual reviews to adjust and finalize the categorization. Naturally, there remain genes without any category assignment.

### 4.6 Annual count of newly researched genes

We counted the genes newly researched in papers and patents each year in the following ways: (*i*) based on the debut year of each gene in papers or patents, whichever earlier for that gene (Fig. 2A); (*ii*) based on the debut year of each gene in papers (Supplementary Fig. S2A in Additional file 1); and (*iii*) based on the debut year of each gene in US, Chinese, or European patents, whichever the earliest for that gene (Supplementary Fig. S2B–D in Additional file 1). Because of the low availability of the patent data after 2020 (Sect. 4.2), we only considered the paper data in the case of (*i*) with the debut year of 2021. Nevertheless, because 2021 is just one year and the paper volume far exceeds the patent volume, the result is unlikely to change much even when the 2021 patent data are considered.

To count the genes related to *Arabidopsis thaliana* or rice among new genes in the category of plant science, we considered the new genes whose paper or patent titles/abstracts matched one of words “Arabidopsis”, “A. thaliana”, and “Arabidopsis thaliana” for *A. thaliana*, and “rice”, “Oryza”, “O. sativa”, and “Oryza sativa” for rice, in a case-insensitive way.

### 4.7 Mathematical modeling of new gene research over time

In our mathematical model, we consider a total of *L* genes and their *M* non-redundant amino acid (AA) sequences across species (i.e., *M* gene copies across species). These sequences can correspond to the UniProtKB/Swiss-Prot entries (Sect. 4.1) and are assumed to have the potential for prospective research. We label these *M* sequences by 1, 2, ⋯, *M* in the temporal order of their debuts in research in papers and patents, and consider all the possible configurations of this labeling with approximately equal *a priori* probabilities. If *X* and *Y* are the number of the member sequences of a given gene and the earliest label of these member sequences, respectively, then the probability of *Y* ≤ *y* for the gene with *X* = *x* is given by

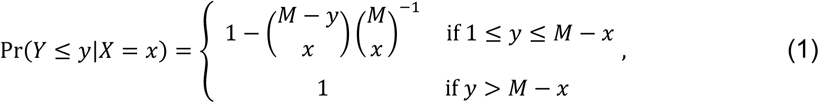

where the binomial coefficient is given by 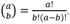 is the number of genes with *x* member sequences each, it satisfies the following relations:

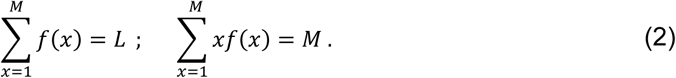

Let *b*(*y*) be the expected number of genes that debuted in research by the time of the *y*^th^-labeled sequence’s debut in research. *b*(*y*) is then expressed as

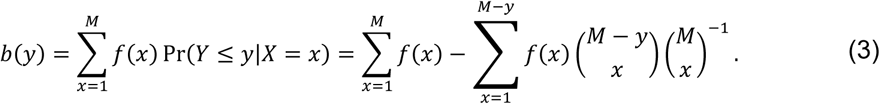

*b*(*y*) in Eq. (3) can be further expressed as the function of time *t*, i.e., *b*(*y*(*t*)) if *y*(*t*) is the total number of the researched sequences by that time *t*. Note that the reporting of a given AA sequence to the scientific community does not necessarily coincide with the research of this sequence. After the reporting of the sequence, the event of its first research may be modeled as a Poisson process with the following *y*(*t*):

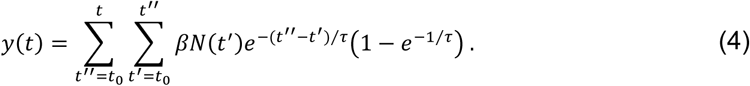

Here, *N*(*t*^’^) is the number of the newly reported, non-redundant sequences in year *t*^’^ obtainable from the UniProtKB/Swiss-Prot entries, *β* is the fraction of the sequences for prospective research (0 ≤ *β* ≤ 1), *t*^’’^ denotes the debut year of each sequence in research, *τ* is the characteristic latent period between the sequence’s first report and research, *t*_0_ is the earliest of year *t*^’^ for the available *N*(*t*^’^) data (i.e., 1939 in the case of the UniProtKB/Swiss-Prot data), and (1 — *e*^−1⁄c^) serves as the normalization factor.

In practice, the reporting years of the sequences in the UniProtKB/Swiss-Prot entries often indicate their first sequencing years, which may not be the same as their actual earliest reporting years, particularly in the case of the early-reported genes before the wide popularization of sequencing technologies. In this regard, we examined the first reporting years in the UniProtKB/Swiss-Prot entries and found some cases before 1997 where each gene’s reporting was spuriously preceded by its debut in research. Therefore, we considered only *t* ≥ 1997 (year) in our modeling.

If there exists *x*^’^ to satisfy 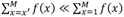, *b*(*y*) in Eq. (3) is simplified as follows:

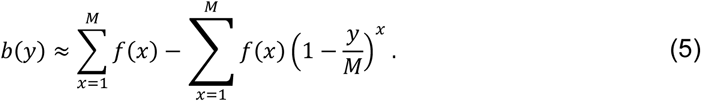

To avoid the overfitting of our model to empirical data, we lump *f*(*x*)’s and reduce their number, as follows:

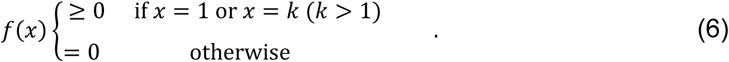

As noted in Sect. 4.1, the non-redundant sequence in each UniProtKB/Swiss-Prot entry is mainly valid at the species level, and therefore genes with *x* = 1 in Eq. (6) correspond to taxonomically-restricted genes (TRGs) or orphans that lack recognizable homologs in other species [39, 40]. Here, we call the other genes taxonomically-dispersed genes (TDGs), and they are lumped to *x* = *k* in Eq. (6) where *k* denotes the effective number of the species for a TDG. Eq. (5) is now converted to

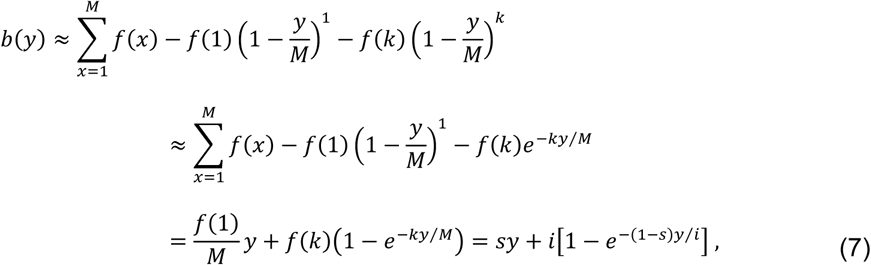

where *s* ≡ *f*(1)/*M*, *i* ≡ *f*(*k*), and we used this relation:

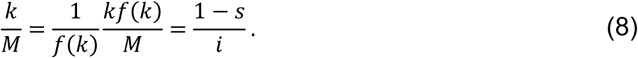

Because the values of *b*(*y*(*t*)) – *b*(*y*(*t –* 1 year)) are available from Fig. 2A, their summation over time leads to *b*(*y*(*t*)) with some constant shift. If we reinterpret *b*(*y*) as this summation of the empirical values, then it is rewritten as

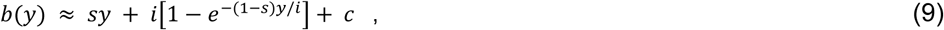

where *c* is a constant.

Next, we seek the parameters in Eq. (9). For each pair of *τ* and *β* in Eq. (4) with (*τ*, *β*)∈ {0, 1, 2, ⋯, 19} ⊗ {0.1, 0.2, 0.3, ⋯,1.0} (τ takes the unit of year), we randomly selected 10^6^ sets of the initial parameters of (*s*, *i*, *c*) in the ranges of (0,1) ⊗ [[, *b*(*t* = 2021)] ⊗ [[, *b*(*t* = 1997)] (*b*(*t*) denotes the above-mentioned empirical data of *b*(*y*(*t*)), *t* takes the unit of year, and we consider only *t* ≥1997 as previously discussed). This random initial parameter selection was conducted by the PCG algorithm in NumPy v1.19.1. Plugging the empirical values of *N*(*t*^’^) in Eq. (4) for *y*(*t*), we minimized the following error functions by the sequential least squares programming (SLSQP in SciPy v1.6.2):

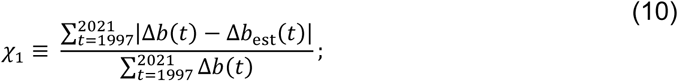

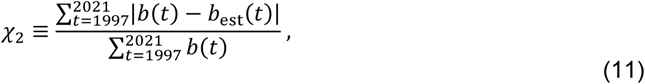

where Δ*b*(*t*) ≡ *b*(*t*) — *b*(*t* — 1 year), Δ*b*_est_(*t*) ≡ *b*_est_(*t*) — *b*_est_(*t* — 1 year), and *b*_est_(*t*) is the right-hand side of Eq. (9) with *y*(*t*) in Eq. (4). The summation in the error functions runs from the year 1997 for the aforementioned reason. Given τ and *β*, the minimization of 3_1_ gives rise to the best-fit *s* and *i*, and the subsequent minimization of 3_2_ with these *s* and *i* gives rise to the best-fit *c*. Fig. 2E presents the heatmap of the resulting 3_1_ and the corresponding graphs of *b*_est_(*t*) and Δ*b*_est_(*t*) as the functions of *y*(*t*) and *t*, respectively. The comparison of *b*_est_(*t*) and *b*(*t*) shows their excellent agreement (3_1_, 3_2_ ≤ 0.07) over a broad range of *τ* and *β*: *P* <10^−4^ for either the resulting 3_1_ or 3_2_ (one-tailed test), from the null distribution of 3_1_ or 3_2_ with the direct use of 10^4^ randomly selected sets of (*s*, *i*, *c*) for each (*τ*, *β*) in the same parameter ranges as above (PCG algorithm in NumPy v1.19.1). We stress that the observed match between *b*_est_(*t*) and *b*(*t*) is not a mere result of any parameter overfitting but reflects the genuine agreement of the empirical data with the model output: *b*(*t*) itself initially curves and approaches a straight line as a function of *y*(*t*) (Fig. 2E) as exactly predicted by Eq. (9) in the model, and thereby always results in 3_2_ < 0.01 whenever 3_1_ ≤ 0.07 is satisfied.

Our model output involves the quantities about TRGs and TDGs mentioned above. First, the fraction of TRG sequences among all the non-redundant sequences of research merit in the model is *f*(1)/*M* = *s*. Across all the *τ* and *β* pairs resulting in 3_1_, 3_2_ ≤ 0.07, we obtained the probability distribution of *s* in Supplementary Fig. S6A (Additional file 1). The expected fraction of the TRGs among all the researched genes as of year *t* is *f*(1)·[*y*(*t*)/*M*] / *b*_est_(*t*) = *sy*(*t*)/ *b*_est_(*t*). In the same 3_1_ and 3_2_ conditions as above, we obtained the probability distribution of the latest available *sy*(*t*) / *b*_est_(*t*) (i.e., *t* = 2021 (year)) as in Supplementary Fig. S6B of Additional file 1.

Because the expected number of TDGs researched as of year *t* is ∼*f*(*k*){1 — [1 — *y*(*t*)/*M*]^k^} ≈ *i*[1 — *e*^−(1–s)y(t)/i^], the expected fraction of the TDGs researched as of year *t*_1_ among the TDGs as of year *t*_2_ (*t*_1_ < *t*_2_) is *F*_TDG_(*t*_1_, *t*_2_) ≡ [1 — *e*^−(1–s)y(t1)/i^]/[1 — *e*^−(1–s)y(t2)/i^]. Because the expected number of TRGs researched as of year *t* is *f*(1)·[*y*(*t*)/*M*] = *sy*(*t*), the expected fraction of the TRGs researched as of year *t*1 among the TRGs as of year *t*_2_ (*t*_1_ < *t*_2_) is *F*_TRG_(*t*_1_, *t*_2_) ≡ *y*(*t*_1_)/ *y*(*t*_2_). In the same 3_1_ and 3_2_ conditions as above, we computed *F*_TDG_(*t*_1_, *t*_2_) and *F*_TRG_(*t*_1_, *t*_2_) when *t*_1_ < 2002 (year) and *t*_2_ = 2021 (year), as presented in Sect. 2.2 for the discussion of the result in Fig. 2A.

### 4.8 Third-generation sequencing trend analysis

We counted papers and patents each year, which were generally related to (*i*) sequencing, or more specifically to (*ii*) third-generation sequencing. Among the papers and patents in the case (*i*), we subsequently obtained the proportion of the case (*ii*) over time. For another analysis, we first collected the papers and patents with specific gene synonyms in their titles/abstracts, counted their cases (*i*) and (*ii*), and then obtained the proportion of the case (*ii*) over time.

Across these analyses, the cases (*i*) and (*ii*) were based on the match of the paper or patent titles/abstracts with the following words: one of “third-generation sequencing”, “long-read sequencing”, “single-molecule sequencing”, “Oxford Nanopore”, “nanopore sequencing”, “GridION”, “PacBio long-read”, “Pac Bio long-read”, “PacBio High-Fidelity”, ”Pac Bio High-Fidelity”, “PacBio HiFi”, ”Pac Bio HiFi”, “HiFi read”, “Pacific Biosciences High-Fidelity”, “Pacific Biosciences HiFi”, “circular consensus sequencing”, “Pacific Biosciences circular consensus sequencing”, “Revio system”, and “Revio sequencer” for the case (*ii*); and one of “sequencing”, “pyrosequencing”, “sequencer”, “DNBSEQ”, “DNBseq”, “DNA-Seq”, “RNA-Seq”, and case-(*ii*) words for the case (*i*). In this matching, the words were treated as case-insensitive, except for “DNBSEQ” and “DNBseq”. We also considered hyphenation variants. Because of the low availability of the patent data after 2020 (Sect. 4.2), we only considered the paper data in the case of 2021. Nevertheless, because 2021 is just one year and the paper volume far exceeds the patent volume, the result is unlikely to change much even when the 2021 patent data are considered.

### 4.9 Identification of gene combinations

To identify the combination of genes that are meaningfully linked in a given paper or patent, we considered the pair, triplet, or entire combination (*n*-tuple with *n* ≥ 2) of the genes that include at least one gene of the main focus in that paper or patent (see Sect. 4.3 for the identification of the genes of the main focus in a paper or patent). As a more conservative approach, we also limited our attention to the pair where both genes are the main focus in the paper or patent; however, this stricter criterion did not alter the overall observed trends in Fig. 3A, and Supplementary Fig. S2E–H and Table S1 in Additional file 1. Because of the low availability of the patent data after 2020 (Sect. 4.2), we only considered the paper data in the case of the gene combinations in 2021. Nevertheless, because 2021 is just one year and the paper volume far exceeds the patent volume, the result is unlikely to change much even when the 2021 patent data are considered.

If any gene in a given pair belongs to a certain thematic gene category, we treated this pair as affiliating with this thematic category (Fig. 3B–D, and Supplementary Fig. S8 in Additional file 1).

For each year or period, we defined the share of a given gene in new pairs as the number of new pairs with this gene divided by the total number of new pairs in that year or period (Fig. 3E–H, and Supplementary Table S1 in Additional file 1). Among the genes with the highest annual shares, Cas9 and GFP are recognized for their utility across disparate domains such as medical and plant sciences. Therefore, for each thematic gene category, we computed the proportion of genes paired with Cas9 or GFP. In this calculation, we focused on the pairings during the early stages of Cas9 and GFP, as these early stages indicate the novelty and innovativeness of the pairings. Specifically, for Cas9 or GFP, we considered the period up to the peak time of its annual paper and patent number.

Additionally, we even considered an earlier stage, the period up to the midpoint of the cumulative paper and patent number at the peak time. As will be seen, the same definitions of the early pairing can apply to other genes of interest beyond Cas9 and GFP. If the annual paper and patent number of such a gene continued to grow until 2021, then 2021 is treated as the peak time in these definitions. Back to the proportion of genes paired with Cas9 or GFP, we considered genes that debuted in papers or patents during similar periods, in order to control for variations from different periods of debut years: one group of genes with debut years of 1990–2000 (Fig. 3I, left and Supplementary Fig. S9A,C in Additional file 1) and the other with 2001–2011 (Fig. 3I, right and Supplementary Fig. S9B,D in Additional file 1). More recent debut years (2012–) were not considered because the number of these genes was relatively small and they may not yet have reached the levels of research stages for the full analysis.

For the statistical significance test of the lower proportion of pharmaceutical genes paired with Cas9 or GFP than that of the whole medical science genes (Fig. 3I), we generated 10^4^ random subsets of the medical science genes and each subset included the same number of genes as pharmaceutical genes. Using these subsets, we obtained the *P* value of the observed proportion of pharmaceutical genes paired with Cas9 or GFP in its above-mentioned early stage (one-tailed test). In addition, we performed the statistical significance test of the relatively low proportion of metabolic engineering (applied biotechnology) genes paired with Cas9 or GFP, compared to those in other categories (Fig. 3I, and Supplementary Fig. S9 in Additional file 1). Specifically, we randomly regrouped genes in major thematic categories—medical science, plant science, metabolic engineering (applied biotechnology), and general microbiology—while keeping the original sizes of these categories. In this way, we generated 10^4^ null configurations of the category membership and obtained the *P* value of the observed proportion of metabolic engineering (applied biotechnology) genes paired with Cas9 or GFP in its above-mentioned early stage and the rank of this proportion among the categories (one-tailed test). All these statistical tests were performed for the genes that debuted in papers or patents during similar periods, as described above.

### 4.10 Definition and analysis of paper and patent falloffs

For each gene, we determined the paper or patent falloff year, which divides its timeline into pre- and post-falloff periods. If annual paper numbers of a given gene are decent enough (≥ 5 papers/year on average for certain three consecutive years), we smoothened the paper time-series with a moving window average (3-year window) to reduce its fluctuations. If the papers in the last window do not exceed a certain threshold (75% of the peak), we considered the latest year of meeting or crossing this threshold after the peak time as the paper falloff year. We defined the patent falloff year in a similar way. Alternations of our methods, however, did not much change our main results.

If annual papers of a given gene are scarce (without any three consecutive years of ≥ 5 papers/year on average), we averaged the paper numbers in year *t*–1, *t*, and *t* +1, and termed this average *X_t_* here. To each year *t*, we assigned one of the following states based on the majority of *X_t_*_–1_, *X_t_*, and *X_t_*_+1_: state I if the majority are ≥1 each; state II if the majority are 0 each; and otherwise, state III. In the case that *t* –1 is the debut year of the gene in papers or patents, or *t* +1 is the last time point, only two of *X_t_*_–1_, *X_t_*, and *X_t_*_+1_ are defined and we assigned one of the following states to year *t*: state I if any of two is ≥1 and both are >0; state II if any of two is 0 and both are <1; and otherwise, state III. In the case that *t* is the debut year of the gene in papers or patents, or *t* is the last time point, only either *X_t_*_–1_ or *X_t_*_+1_ is defined and we assigned one of the following states to year *t*: state I if the value is ≥1; state II if the value is 0; and otherwise, state III. If the paper time-series includes state I sometime before the last point and state II or III at the last point, we considered the year next to the latest time of state I as the paper falloff year. We defined the patent falloff year in a similar way. Alternations of our methods, however, did not much change our main results.

To investigate paper and patent falloff patterns, we obtained the proportion of genes whose paper falloffs preceded the patent falloffs and vice versa (Fig. 4B). In this calculation, we used the same period (–2020) for both paper and patent time-series for their fair comparison and considered only the genes that satisfy the following conditions: (*i*) any of the paper and patent time-series includes the falloff year defined above; and (*ii*) neither the paper nor patent time-series is fully spanned by states II and III defined above. In addition, we considered the genes that debuted in papers or patents during similar periods (1990– 2000 and 2001–2011, but not 2012–) for the same reason in Sect. 4.9.

To examine whether the policy requirement of non-overlap of a patent content with prior art is responsible for the observed tendency of the paper and patent falloffs (Fig. 4B), we filtered out papers and patents with highly similar contents to their prior works and re-analyzed the data (Supplementary Fig. S10 in Additional file 1). This test ensures a fair paper and patent treatment, with the analyzed papers being as dissimilar to their prior works as the patents are. Specifically, we used PaECTER, a citation-informed sentence transformer [98] and obtained the cosine similarity of every pair of papers (patents) for each gene, based on the paper (patent) titles and abstracts. If the total number of papers (patents) exceeded 5,000 for a given gene, we only used 5,000 randomly selected papers (patents) for the similarity calculation, which would otherwise become computationally impractical. For the papers (patents) of each gene in a particular year, we filtered out the later papers (patents) with the similarity at or above a certain threshold, and repeated this process across all the considered years in chronological order. The similarity threshold here (0.97) was determined by the manual inspection of the contents of paper or patent pairs with varying threshold values. After the filtering, we obtained the proportion of genes whose paper falloffs preceded the patent falloffs and vice versa. Regarding the control group without the filtering, we used the aforementioned 5,000 papers (patents) for each gene if this gene occurred in more than 5,000 papers (patents), and otherwise, all its papers (patents).

For the statistical significance test of the dominance of genes whose patent falloffs preceded the paper falloffs (Fig. 4B, and Supplementary Fig. S10 in Additional file 1), we randomly swapped the paper and patent time-series of each gene and generated 10^4^ null configurations. Using these null configurations, we obtained the *P* value of the observed proportion of genes whose patent falloffs preceded the paper falloffs (one-tailed test). This statistical test was performed for the genes that debuted in papers or patents during similar periods (Sect. 4.9).

### 4.11 The total paper versus patent numbers of each gene

The practical upper limit of the patent number of a gene can be expressed as a function of its paper number, *y* = *αx^γ^* where *x* and *y* are the paper and patent numbers, respectively (Supplementary Fig. S11 in Additional file 1). The parameters *γ* and *α* were initially found by the fitting of the function with the least square method (polyfit in NumPy v1.19.1) and then adjusted manually. To test the statistical significance of the observed fraction of the genes below or on the upper limit, we randomly permuted the paper numbers of the genes (or equivalently, their patent numbers) and obtained the probability of the fraction of the genes (below or on the same above-mentioned upper limit) being greater than or equal to the observed fraction. This probability serves as the *P* value of the observed fraction (one-tailed test) and was computed with 10^4^ trials of the randomization by the PCG algorithm in NumPy v1.19.1.

### 4.12 Selection of genes that signify biotechnology innovation

We first calculated the share of each gene in new annual pairs of genes, as defined in Sect. 4.9. We then identified the genes of the highest shares for ≥5 consecutive years or among the top five share genes for ≥10 years in 1990–2021. They were a total of nine genes that signify biotechnology innovation: Cas9 for genome editing [42], PD-L1 for immunomodulation and immunotherapy [56], mTOR, phosphatidylinositol 3-kinase (PI3-K), and TP53 mainly for cancer research [57, 58, 99], GFP and luciferase for live cell imaging [59, 60], β-catenin mainly for regenerative medicine and cancer research [61, 62], and β-galactosidase for DNA cloning [41]. We excluded PI3-K from further analysis due to its strong contextual overlap with mTOR through the biochemical pathway sharing. The remaining eight genes are referred to as *iconic genes* in this study. The results of our iconic-gene analysis, presented in Supplementary Fig. S12–S14 (Sect. 4.15, 4.16), remain qualitatively similar when we instead analyze non-iconic but still high-share genes (Supplementary Fig. S15 and Table S1; see Additional file 1).

### 4.13 Clinical trial analysis

We sought the papers with a given gene as the main focus (Sect. 4.3), while each paper contains the titles/abstracts with “clinicaltrials.gov” (case-insensitive) and “NCT” followed by an eight-digit number. Using this National Clinical Trials (NCT) number, we identified the type of the sponsor for that clinical trial from ClinicalTrials.gov (June 6, 2024), managed by the U.S. National Library of Medicine as a registry of global clinical-trial activities. Among the sponsor types, we only considered “INDUSTRY”, “NIH”, “OTHER_GOV”, “FED”, and “OTHER”. In contrast to “INDUSTRY”, we confirmed the public nature of the rest sponsor types by reviewing their specific sponsors.

Of the above “OTHER” non-industry sponsors, we classified a subset as “hospital” with each name containing “hospital”, “medical center”, “cancer center”, “health center”, “health centre”, “health network”, “rigshospitalet”, “hôpitaux”, or “centre hospitalier”. Of the remaining “OTHER” sponsors, we classified a subset as “university” with each name containing “university”, “college”, “academy”, “academia”, “academic”, “universitaire”, “universidad”, “universidade”, “universita”, “universität”, “universiteit”, “school of medicine”, “medical school”, “college of medicine”, or “faculty of medicine”. The rest non-industry sponsors were classified as “miscellany”.

Since September 2007, the Food and Drug Administration Amendments Act of 2007 has mandated the posting of trials to ClinicalTrials.gov. According to our observation, it typically took ∼6 years between the debut of a gene in papers/patents and its first trial, suggesting 2002– as the appropriate debut years of genes for the analysis. In addition, the genes of relatively recent debut years (2012–) were not considered because the number of these genes was relatively small and they may not yet have reached the levels of research stages for the full analysis. As a result, we focused on the genes of the debut years 2002–2011.

Among these genes in clinical-trial papers as their main focus, we obtained the proportion of the genes in those papers before the patent falloffs (Fig. 4C, left “Pre-falloff”), and the proportion after (or at) the patent falloffs or with the patent time-series fully spanned by states II and III defined in Sect. 4.10 (Fig. 4C, left “Post-falloff”). To calculate the probability of industry (or other) sponsorship for a clinical trial with each gene before its patent falloff, we obtained the average proportion of clinical trials with industry (or other) sponsorship described above, over all the genes in the clinical-trial papers before the patent falloffs (Fig. 4C, right “Pre-falloff” and Supplementary Fig. S16A,B in Additional file 1).

Similarly, we calculated the probability of industry (or other) sponsorship for a clinical trial with each gene after (or at) its patent falloff or of the patent time-series fully spanned by states II and III defined in Sect. 4.10 (Fig. 4C, right “Post-falloff” and Supplementary Fig. S16A,B in Additional file 1).

For the statistical significance test of the high probability of non-industry sponsorship for a clinical trial with each gene compared to industry sponsorship, we randomly but equally reassigned industry or other sponsorship to each clinical trial and generated the 10^4^ null configurations. From these null configurations, we obtained the *P* value of the observed probability of non-industry sponsorship in the case before the patent falloff, or in the case after (or at) the patent falloff or of the patent time-series fully spanned by states II and III (one-tailed test).

Alternatively, rather than starting with clinical-trial papers as above, we repeated the whole analysis by directly screening the XML file of ClinicalTrials.gov for trial-related genes and data (Supplementary Table S4 in Additional file 1). The gene search here was performed by the string-matching algorithm in Sect. 4.3. Still, the main findings in Fig. 4C remained valid under this re-analysis (Supplementary Fig. S16C,D in Additional file 1).

### 4.14 Patent citation analysis

We collected the data of patent citations of papers from SciSciNet [85]. For compatibility with our paper data, the PubMed reference numbers (PMIDs) of papers in SciSciNet were retrieved from OpenAlex (May 25, 2024) [84]. We focused on highly patent-cited works, the papers that received ≥3 patent citations each within the first three years of publications.

However, alternative criteria of high patent citations did not qualitatively change our analysis result. Among the genes of the main focus in the highly patent-cited works, we obtained the proportion of genes in those works before the patent falloffs (Fig. 4D, “Pre-falloff”), and the proportion after (or at) the patent falloffs or with the patent time-series fully spanned by states II and III defined in Sect. 4.10 (Fig. 4D, “Post-falloff”). To control for variations from different debut years of genes, we considered the genes of similar debut periods, 1990– 2000 and 2001–2011 separately. More recent debut years (2012–) were not considered for the same reason in Sect. 4.9.

### 4.15 Analysis of the teaming with new iconic genes

We considered genes that had the patent falloff years (defined in Sect. 4.10) and were paired with any iconic genes until the peak times (Sect. 4.9) of these iconic genes. Among them, we obtained the proportion paired with ≥1, ≥2, or ≥3 new iconic genes after (or at) the patent falloffs, until the peak times of the iconic genes (Fig. 5B). In addition, for a given iconic gene, we considered genes paired with this iconic gene until its peak time. Among them, we obtained the proportion of genes initially paired with that early iconic gene before/after the patent falloffs (Fig. 5B, and Supplementary Fig. S12A in Additional file 1).

Similarly, we obtained the proportion for the midpoint of the peak time as defined in Sect. 4.9, instead of the peak time (Supplementary Fig. S12B in Additional file 1). All these analyses were performed for the genes that debuted in papers or patents during a similar period, 1990–2000. Genes with post-2000 debuts were not considered due to the fundamentally low overlap of their active research periods with the early stages of the iconic genes.

For a given iconic gene, we next focused on the genes that satisfy the following conditions: (*i*) they were initially paired with this iconic gene until its peak time or the midpoint of the peak time, after (or at) the patent falloffs; (*ii*) their paper time-series are not fully spanned by states II and III defined in Sect. 4.10; (*iii*) the patent falloffs preceded or coincided with the paper falloffs. Among these genes, we obtained the proportion of genes initially paired with that early iconic gene before/after the paper falloffs (Fig. 5C, and Supplementary Fig. S13 in Additional file 1). In addition, we performed the statistical significance test of the higher proportion of those genes before the paper falloffs than after (or at) the paper falloffs, observed for most iconic genes. Specifically, we randomly reassigned the initial pairing times of each iconic gene with equal *a priori* probabilities over the range from the iconic gene’s debut year or its paired gene’s patent falloff year (whichever later) to the latest point of at least one paper or patent of this paired gene. Across all the relevant iconic genes and their paired genes, we generated 10^4^ null configurations of the initial pairing times and obtained the *P* value of the observed, joint proportions of genes initially paired with the iconic genes before the paper falloffs. All the analyses were performed for the genes that debuted in papers or patents during a similar period (1990– 2000), as discussed above.

### 4.16 Hypothetical scenario of exploratory-opportunity loss

We quantified how much the exploratory opportunities of genetic research depend on persistent scientific activity beyond short industrial drive. For this quantification, we conducted a “thought experiment” for each iconic gene, as follows.

Define set *A* as the genes that teamed with this iconic gene until its peak time and satisfy the following condition (*i*) or (*ii*): (*i*) they were initially paired with that early iconic gene after (or at) the patent falloffs and the patent falloffs preceded or coincided with the paper falloffs; or (*ii*) the patent time-series are fully spanned by states II and III (defined in Sect. 4.10) but the paper time-series are not. In addition, define set *B* as a subset of *A*, where genes were initially paired with that early iconic gene before the paper falloffs. Also, define set *C* as a subset of *A*, where genes were initially paired with that early iconic gene after (or at) the paper falloffs.

For the genes initially paired with a given early iconic gene with their limited patent volumes (i.e., set *A*), we consider a hypothetical scenario that the papers undergo immediate falloffs due to the limited patent volumes. This scenario allows us to estimate the maximum extent to which the early teaming with this iconic gene would be lost if the research activities become hindered in step with the industry’s limited interest. The estimated proportion of genes that would lose this early teaming is given by (|*A*| — *x*_0_)⁄|*A*| (Fig. 5D, and Supplementary Fig. S14 in Additional file 1), where *x*_0_ is the number of genes that would still maintain the teaming despite the paper falloffs. Here, *x*_0_ ≡ *P*|*B*| + |*C*| with *P* ≡ |*C*|⁄|*A*|. *P* denotes the conditional probability that each gene in *B* would still maintain the teaming.

By definition, all genes in *C* would clearly maintain the teaming.

We further define set *D* as all genes that teamed with a given early iconic gene. We also define set *E* as *E* ≡ *D* –*B*. Among all the genes in *D* whether affected or not by the above hypothetical scenario, the estimated proportion of genes that would lose the early teaming with the iconic gene is given by (|*D*| — *x*_1_)⁄|*D*| (Fig. 5D, and Supplementary Fig. S14 in Additional file 1), where *x*_1_ ≡ *P*|*B*| + |*E*| with *P* defined previously. This derivation is similar to the above formulation.

Additionally, we conducted the same thought experiment, but for the teaming with each iconic gene until the midpoint of its peak time as defined in Sect. 4.9. All these analyses were performed for the genes that debuted in papers or patents during a similar period (1990–2000), as discussed in Sect. 4.15. In the example of COX-2 or PDE5, the estimated proportion of the chances to lose the early teaming with a particular iconic gene is given by 1 — *P*, calculated for this iconic gene.

## Availability of data and materials

All relevant data are included in this article and Additional files 1–7. Source codes are available in the public repository GitHub, https://github.com/WooJoongKim0107/GeneBib_demo.

## Acknowledgements

We thank Taekyun Kim, Hong Lok Lung, Seolmin Yang, Daehyun Kim, Liming Xiong, Yiji Xia, Adam G. Craig, and Anna Oi Wah Leung for useful discussions. This work was supported by the National Research Foundation of Korea Grants RS-2023-00263411 and RS-2024-00353098 funded by the Ministry of Science and ICT (J.C., W.J.K., R.L., and C.-M.G.), Hong Kong Baptist University Start-up Grant for New Academics (R.L. and P.-J.K.), the National Science Foundation Grant Award EF-2133863 and the National Research Foundation of Korea, Global Humanities and Social Sciences Convergence Research Program 2024S1A5C3A02042671 (H.Y.). We also acknowledge the support of the UNIST Supercomputing Center for the computing resources.

## References

1. Ahmed N, Wahed M, Thompson NC. The growing influence of industry in AI research. Science. 2023;379(6635): 884–6.

2. Kumar A, Blum J, Le TT, Havelange N, Magini D, Yoon IK. The mRNA vaccine development landscape for infectious diseases. Nat Rev Drug Discov. 2022;21(5): 333–4.

3. Gibney E. Quantum gold rush: the private funding pouring into quantum start-ups. Nature. 2019;574: 22–4.

4. March JG. Exploration and exploitation in organizational learning. Organ Sci. 1991;2(1): 71–87.

5. Christensen CM. The Innovator’s Dilemma: When New Technologies Cause Great Firms to Fail. Boston, MA: Harvard Business School Press; 1997.

6. Agarwal R, Gaule P. What drives innovation? Lessons from COVID-19 R&D. J Health Econ. 2022;82: 102591.

7. Krieger J, Li D, Papanikolaou D. Missing novelty in drug development. Rev Financ Stud. 2022;35: 636–79.

8. Teece DJ. Profiting from technological innovation: Implications for integration, collaboration, licensing and public policy. Res Policy. 1986;15: 285–305.

9. Blumenthal D, Causino N, Campbell E, Louis KS. Relationships between academic institutions and industry in the life sciences—an industry survey. N Engl J Med. 1996;334(6): 368–73.

10. Pellegrino G. Barriers to innovation in young and mature firms. J Evol Econ. 2018;28: 181–206.

11. Pisano G. Profiting from innovation and the intellectual property revolution. Res Policy. 2006;35(8): 1122–30.

12. Georgescu I. Bringing back the golden days of Bell Labs. Nat Rev Phys. 2022;4(2):76–8.

13. Shi F, Evans J. Surprising combinations of research contents and contexts are related to impact and emerge with scientific outsiders from distant disciplines. Nat Commun. 2023;14(1): 1641.

14. Youn H, Strumsky D, Bettencourt LMA, Lobo J. Invention as a combinatorial process: evidence from US patents. J R Soc Interface. 2015;12(106): 20150272.

15. Fink TMA, Reeves M. How much can we influence the rate of innovation? Sci Adv. 2019;5(1): eaat6107.

16. Ke Q. Technological impact of biomedical research: The role of basicness and novelty. Res Policy. 2020;49: 104071.

17. Lee ED, Kempes CP, West GB. Idea engines: Unifying innovation & obsolescence from markets & genetic evolution to science. Proc Natl Acad Sci U S A. 2024;121(6): e2312468120.

18. Lee ED, Kempes CP, Laubichler MD, Hamilton MJ, Lockhart JW, Neffke F, et al. Synthesis of innovation and obsolescence. arXiv:2505.05182 [Preprint]. 2025 [cited 2026 July 9]. Available from: https://arxiv.org/abs/2505.05182

19. Ahmadpoor M, Jones BF. The dual frontier: Patented inventions and prior scientific advance. Science. 2017;357(6351): 583–7.

20. Poege F, Harhoff D, Gaessler F, Baruffaldi S. Science quality and the value of inventions. Sci Adv. 2019;5(12): eaay7323.

21. Kwon S. Underappreciated government research support in patents. Science. 2024; 385(6712): 936–8.

22. Park M, Leahey E, Funk RJ. Papers and patents are becoming less disruptive over time. Nature. 2023;613(7942): 138–44.

23. Bloom N, Jones CI, Van Reenen J, Webb M. Are ideas getting harder to find? Am Econ Rev. 2020;110: 1104–44.

24. Gates AJ, Gysi DM, Kellis M, Barabási A-L. A wealth of discovery built on the Human Genome Project - by the numbers. Nature. 2021;590(7845): 212–5.

25. McElheny VK. Drawing the Map of Life: Inside the Human Genome Project. New York, NY: Basic Books; 2010.

26. McGuire AL, Gabriel S, Tishkoff SA, Wonkam A, Chakravarti A, Furlong EEM, et al. The road ahead in genetics and genomics. Nat Rev Genet. 2020;21(10): 581–96.

27. Wu W, Zhang HH, J. WM, B KP. Gene Biotechnology. 2nd ed. Boca Raton, FL: CRC Press; 2003.

28. Stoeger T, Amaral LAN. The characteristics of early-stage research into human genes are substantially different from subsequent research. PLOS Biol. 2022;20(1): e3001520.

29. Hoffmann M, Kleine-Weber H, Schroeder S, Kruger N, Herrler T, Erichsen S, et al. SARS-CoV-2 cell entry depends on ACE2 and TMPRSS2 and is blocked by a clinically proven protease inhibitor. Cell. 2020;181(2): 271–80.

30. Stoeger T, Amaral LAN. Meta-Research: COVID-19 research risks ignoring important host genes due to pre-established research patterns. eLife. 2020;9: e61981.

31. The UniProt Consortium. UniProt: the universal protein knowledgebase in 2021. Nucleic Acids Res. 2021;49(D1): D480–9.

32. Narin F, Noma E. Is technology becoming science? Scientometrics. 1985;7: 369–81.

33. Narin F, Noma E, Perry R. Patents as indicators of corporate technological strength. Res Policy. 1987;16: 143–55.

34. Meyer M. Does science push technology? Patents citing scientific literature. Res Policy. 2000;29: 409–34.

35. Sampat B, Williams HL. How do patents affect follow-on innovation? Evidence from the human genome. Am Econ Rev. 2019;109(1): 203–36.

36. Stephanopoulos G, Aristidou AA, Nielsen JH. Metabolic Engineering: Principles and Methodologies. San Diego: Academic Press; 1998.

37. Sun L, Lee JW, Yook S, Lane S, Sun ZQ, Kim SR, et al. Complete and efficient conversion of plant cell wall hemicellulose into high-value bioproducts by engineered yeast. Nat Commun. 2021;12(1): 4975.

38. Provart NJ, Alonso J, Assmann SM, Bergmann D, Brady SM, Brkljacic J, et al. 50 years of *Arabidopsis* research: highlights and future directions. New Phytol. 2016;209(3): 921–44.

39. Wilson GA, Bertrand N, Patel Y, Hughes JB, Feil EJ, Field D. Orphans as taxonomically restricted and ecologically important genes. Microbiol-Sgm. 2005;151: 2499–501.

40. Khalturin K, Hemmrich G, Fraune S, Augustin R, Bosch TCG. More than just orphans: are taxonomically-restricted genes important in evolution? Trends Genet. 2009;25(9): 404–13.

41. Ullmann A, Jacob F, Monod J. Characterization by *in vitro* complementation of a peptide corresponding to an operator-proximal segment of beta-galactosidase structural gene of *Escherichia coli*. J Mol Biol. 1967;24(2): 339–43.

42. Jinek M, Chylinski K, Fonfara I, Hauer M, Doudna JA, Charpentier E. A programmable dual-RNA-guided DNA endonuclease in adaptive bacterial immunity. Science. 2012;337(6096): 816–21.

43. Topol EJ. Failing the public health—rofecoxib, Merck, and the FDA. N Engl J Med. 2004;351: 1707–9.

44. Shin DM, et al. Chemoprevention of head and neck cancer by simultaneous blocking of epidermal growth factor receptor and cyclooxygenase-2 signaling pathways: preclinical and clinical studies. Clin Cancer Res. 2013;19(5): 1244–56.

45. Saba NF, et al. Chemoprevention of head and neck cancer with celecoxib and erlotinib: results of a phase Ib and pharmacokinetic study. Cancer Prev Res. 2014;7(3): 283–91.

46. Groeger AL, et al. Cyclooxygenase-2 generates anti-inflammatory mediators from omega-3 fatty acids. Nat Chem Biol. 2010;6(6): 433–41.

47. Chen JS, et al. Nonsteroidal anti-inflammatory drugs dampen the cytokine and antibody response to SARS-CoV-2 infection. J virol. 2021;95(7): e00014–21.

48. Jackson G, Gillies H, Osterloh I. Past, present, and future: a 7-year update of Viagra^®^ (sildenafil citrate). Int J Clin Pract 2005;59(6): 680–91.

49. Bhogal S, Khraisha O, Al Madani M, Treece J, Baumrucker SJ, Paul TK. Sildenafil for pulmonary arterial hypertension. Am J Ther. 2019;26(4): e520–6.

50. U.S. National Institutes of Health. ClinicalTrials.gov. Available from: http://www.clinicaltrials.gov.

51. Tinsley HN, et al. Colon tumor cell growth-inhibitory activity of sulindac sulfide and other nonsteroidal anti-inflammatory drugs is associated with phosphodiesterase 5 inhibition.” Cancer Prev Res. 2010;3(10): 1303–13.

52. Tinsley HN, et al. Inhibition of PDE5 by sulindac sulfide selectively induces apoptosis and attenuates oncogenic Wnt/β-catenin–mediated transcription in human breast tumor cells. Cancer Prev Res. 2011;4(8): 1275–84.

53. Nelson SF, Miceli MC, Elashoff RM, Sweeney HL, Victor RG. PDE5 inhibition alleviates functional muscle ischemia in boys with Duchenne muscular dystrophy. Neurology. 2014;82(23): 2085–91.

54. Wang R, Lei X, Rakesh CK. PDE5 inhibitor tadalafil and hydroxychloroquine cotreatment provides synergistic protection against type 2 diabetes and myocardial infarction in mice. J Pharmacol Exp Ther. 2017;361(1): 29–38.

55. Amedro P, et al. Efficacy of phosphodiesterase type 5 inhibitors in univentricular congenital heart disease: the SV-INHIBITION study design. ESC Heart Fail. 2020;7(2): 747–56.

56. Ohaegbulam KC, Assal A, Lazar-Molnar E, Yao Y, Zang XX. Human cancer immunotherapy with antibodies to the PD-1 and PD-L1 pathway. Trends Mol Med. 2015;21(1): 24–33.

57. Zou ZL, Tao T, Li HM, Zhu X. mTOR signaling pathway and mTOR inhibitors in cancer: progress and challenges. Cell Biosci. 2020;10(1): 31.

58. Mantovani F, Collavin L, Del Sal G. Mutant p53 as a guardian of the cancer cell. Cell Death Differ. 2019;26(2): 199–212.

59. Lippincott-Schwartz J, Patterson GH. Development and use of fluorescent protein markers in living cells. Science. 2003;300(5616): 87-91.

60. Liu S, Su YC, Lin MZ, Ronald JA. Brightening up biology: advances in luciferase systems for *in vivo* Imaging. Acs Chem Biol. 2021;16(12): 2707–18.

61. Liu JQ, Xiao Q, Xiao JN, Niu CX, Li YY, Zhang XJ, et al. Wnt/beta-catenin signalling: function, biological mechanisms, and therapeutic opportunities. Signal Transduct Tar. 2022;7(1): 3.

62. Zhang Y, Wang X. Targeting the Wnt/beta-catenin signaling pathway in cancer. J Hematol Oncol. 2020;13(1): 165.

63. English K, Barry FP, Field-Corbett CP, Mahon BP. IFN-γ and TNF-α differentially regulate immunomodulation by murine mesenchymal stem cells. Immunol Lett. 2007;110(2): 91–100.

64. Liu X, Ji Q, Zhang C, Liu X, Liu Y, Liu N, et al. miR-30a acts as a tumor suppressor by double-targeting COX-2 and BCL9 in H. pylori gastric cancer models. Sci Rep. 2017;7(1): 7113.

65. Biswas KH, Sopory S, Visweswariah SS. The GAF domain of the cGMP-binding, cGMP-specific phosphodiesterase (PDE5) is a sensor and a sink for cGMP. Biochemistry. 2008;47(11): 3534–43.

66. Küçük A, Yucel M, Erkasap N, Tosun M, Koken T, Ozkurt M, et al. The effects of PDE5 inhibitory drugs on renal ischemia/reperfusion injury in rats. Mol Biol Rep. 2012;(39): 9775–82.

67. Tavallai M, Hamed HA, Roberts JL, Cruickshanks N, Chuckalovcak J, Poklepovic A, et al. Nexavar/Stivarga and viagra interact to kill tumor cells. J Cell Physiol. 2015;230(9): 2281–98.

68. Krimsky S. Science in the Private Interest: Has the Lure of Profits Corrupted Biomedical Research? Lanham, MD: Rowman & Littlefield; 2003.

69. BP cuts funding for ‘most promising’ biofuel. Investigate Midwest. August 20, 2015; https://investigatemidwest.org/2015/08/20/bp-cuts-funding-for-most-promising-biofuel-2.

70. Stocker M, Baffes J, Vorisek D. What triggered the oil price plunge of 2014–2016 and why it failed to deliver an economic impetus in eight charts. World Bank Blogs. January 18, 2018; https://blogs.worldbank.org/en/developmenttalk/what-triggered-oil-price-plunge-2014-2016-and-why-it-failed-deliver-economic-impetus-eight-charts.

71. Asase RV, Okechukwu QN, Ivantsova MN. Biofuels: present and future. Environ Dev Sustain. 2024; 10.1007/s10668-024-04992-w.

72. Mandt R, Seetharam K, Cheng CHM. Federal R&D funding: the bedrock of national innovation. MIT Sci Policy Rev. 2020;1: 44–54.

73. Yun J, Kim P-J, Jeong H. Anatomy of scientific evolution. PLOS ONE. 2015;10(2): e0117388.

74. Bush V. Science the Endless Frontier: A Report to the President. Washington, DC: United States Office of Scientific Research and Development; 1945.

75. Stokes DE. Pasteur’s Quadrant: Basic Science and Technological Innovation. Washington, DC: Brookings Institution; 1997.

76. Mokyr J. The Gifts of Athena: The Origin of the Knowledge Economy. Princeton, NJ: Princeton University; 2002.

77. Perkmann M, Tartari V, McKelvey M, Autio E, Broström A, D’Este P, et al. Academic engagement and commercialisation: A review of the literature on university–industry relations. Res Policy. 2013;42(2): 423–42.

78. Hiltzik MA. Dealers of Lightning: Xerox PARC and the Dawn of the Computer Age. New York, NY: HarperCollins; 1999.

79. Haessler P, Giones F, Brem A. The who and how of commercializing emerging technologies: A technology-focused review. Technovation. 2023;121: 102637.

80. Levinthal DA. The slow pace of rapid technological change: Gradualism and punctuation in technological change. Ind Corp Change. 1998;7(2): 217–47.

81. Scaringella L, Miles RE, Truong Y. Customers involvement and firm absorptive capacity in radical innovation: The case of technological spin-offs. Technol Forecast Soc Change. 2017;120: 144–62.

82. Arthur WB. The structure of invention. Res Policy 2007;36: 274–87.

83. Marx M, Fuegi A. Reliance on science by inventors: Hybrid extraction of in-text patent-to-article citations. J Econ Manage Strat. 2022;31(2): 369–92.

84. Priem J, Piwowar H, Orr R. OpenAlex: A fully-open index of scholarly works, authors, venues, institutions, and concepts. arXiv:2205.01833 [Preprint]. 2022 [cited 2025 May 2]. Available from: https://arxiv.org/abs/2205.01833

85. Lin Z, Yin Y, Liu L, Wang D. SciSciNet: A large-scale open data lake for the science of science research. Sci Data. 2023;10(1): 315.

86. Hocher A, Laursen SP, Radford P, Tyson J, Lambert C, Stevens KM, et al. Histones with an unconventional DNA-binding mode *in vitro* are major chromatin constituents in the bacterium *Bdellovibrio bacteriovorus*. Nat Microbiol. 2023; 8: 2006–19.

87. Arnold C. Death by climate change. Nat Clim Change. 2022;12(7):607–9.

88. Perry BL, Aronson B, Pescosolido BA. Pandemic precarity: COVID-19 is exposing and exacerbating inequalities in the American heartland. Proc Natl Acad Sci U S A. 2021;118(8): e2020685118.

89. Chen CY, Kahanamoku SS, Tripati A, Alegado RA, Morris VR, Andrade K, et al. Systemic racial disparities in funding rates at the National Science Foundation. eLife. 2022;11: e83071.

90. Bairoch A, Apweiler R, Wu CH, Barker WC, Boeckmann B, Ferro S, et al. The universal protein resource (UniProt). Nucleic Acids Res. 2005;33: D154–D9.

91. Browet A, Absil P-A, Dooren PV. Fast community detection using local neighbourhood search. arXiv:1308.6276 [Preprint]. 2013 [cited 2025 May 2]. Available from: https://arxiv.org/abs/1308.6276

92. Lin Y, Michel J-B, Aiden EL, Orwant J, Brockman W, Petrov S. Syntactic annotations for the Google Books Ngram Corpus. Proceedings of the 50th Annual Meeting of the Association for Computational Linguistics. 2012: 169–74.

93. Sayers EW, Bolton EE, Brister JR, Canese K, Chan J, Comeau DC, et al. Database resources of the National Center for Biotechnology Information in 2023. Nucleic Acids Res. 2022;50: D20–6.

94. Fredkin E. Trie Memory. Commun Acm. 1960;3(9): 490–9.

95. Wei CH, Allot A, Lai PT, Leaman R, Tian S, Luo L, et al. PubTator 3.0: an AI-powered literature resource for unlocking biomedical knowledge. Nucleic Acids Res, 2024;52(W1): W540–6.

96. The Gene Ontology Consortium. The Gene Ontology resource: enriching a GOld mine. Nucleic Acids Res. 2021;49(D1): D325–34.

97. Wishart DS, Feunang YD, Guo AC, Lo EJ, Marcu A, Grant JR, et al. DrugBank 5.0: a major update to the DrugBank database for 2018. Nucleic Acids Res. 2018;46(D1): D1074–82.

98. Ghosh M, Erhardt S, Rose ME, Buunk E, Harhoff D. PaECTER: patent-level representation learning using citation-informed transformers. arXiv:2402.19411 [Preprint]. 2024 [cited 2025 May 2]. Available from: https://arxiv.org/abs/2402.19411

99. Fruman DA, Christian R. PI3K and cancer: lessons, challenges and opportunities. Nat Rev Drug Discov. 2014;13(2): 140–56.

