## Additional files 1 to 7 for "Scientific exploration outlasts industrial momentum and sustains innovation in genetic research: survey of 20 million papers and patents": Additional_File_1.pdf

### Supplementary Figures and Tables

Junghun Chae*, WooJoong Kim*, Woochul Jung*, Dawoon Jeong*, Roktaek Lim*, Manoj Chamlagain, Giju Jung, Juneil Jang, Jae Won Lee, Nam Kyu Kang, Kwangryul Baek, Jonghyeok Shin, Ye-Gi Lee, Hyun Gi Koh, Chanwoo Kim, Sangdo Yook, Allen Ka Loon Cheung, Yong-Su Jin, Hyejin Youn†, Pan-Jun Kim†, and Cheol-Min Ghim†

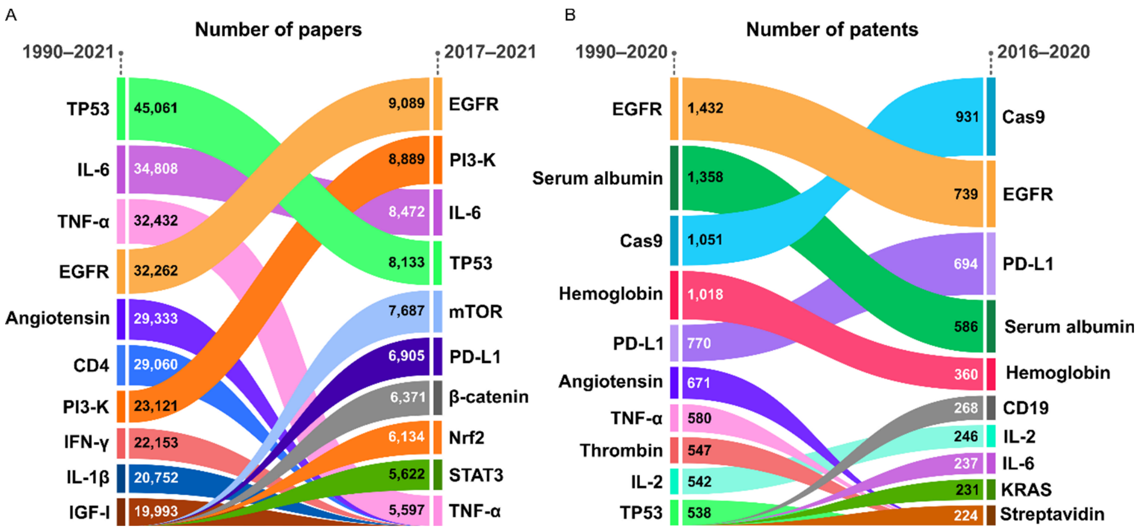


**Supplementary Fig. S1. Genes of frequent research focus.** Related to Fig. 1E,F. Ten most frequent genes of the main focus in papers (**A**) or patents (**B**) of all time (1990–2021 for papers and 1990–2020 for patents; left side) and latest five years (2017–2021 for papers and 2016–2020 for patents; right side). For the identification of the genes of the main focus in papers or patents, o Sect. 4.3. The genes are vertically ordered by their relevant paper or patent number in a given period. Each gene is indicated by a ribbon with a vertical width, proportional to the paper or patent number.

*These authors contributed equally.

†Correspondence and requests for materials should be addressed to
H.Y., P.-J.K., or C.-M.G..

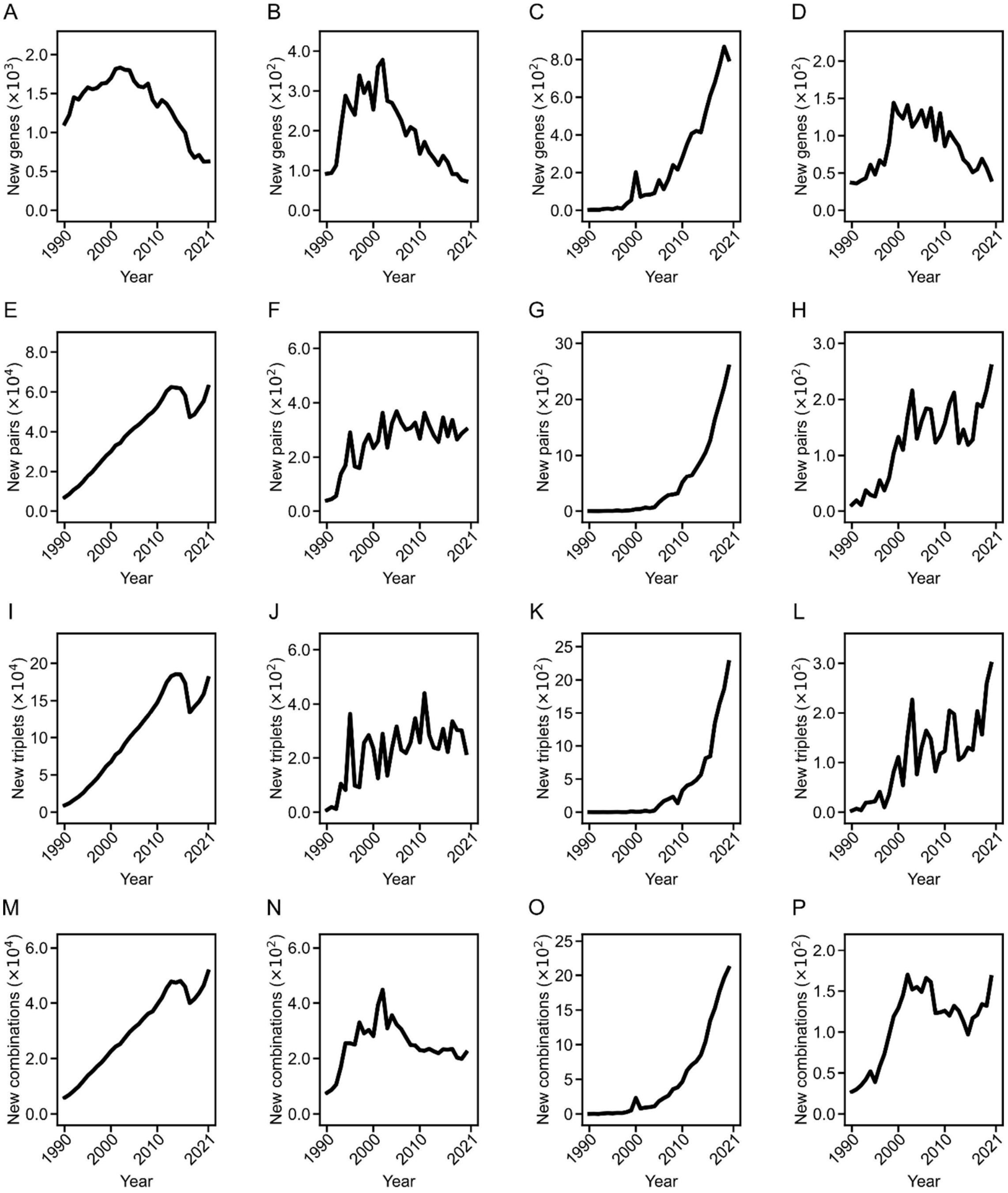


**Supplementary Fig. S2. Annual counts of newly researched genes and novel gene combinations.** Related to Fig. 2A, 3A. **A–D**, Annual counts of new genes in only papers (**A**), and only patents from the US (**B**), China (**C**), or Europe (**D**) whichever the earliest. **E–P**, Annual counts of new pairs (**E–H**), new triplets (**I–L**), or new entire combinations (**M–P**) of genes in only papers (**E**, **I**, and **M**), and only patents from the US (**F**, **J**, and **N**), China (**G**, **K**, and **O**), or Europe (**H**, **L**, and **P**) whichever the earliest.

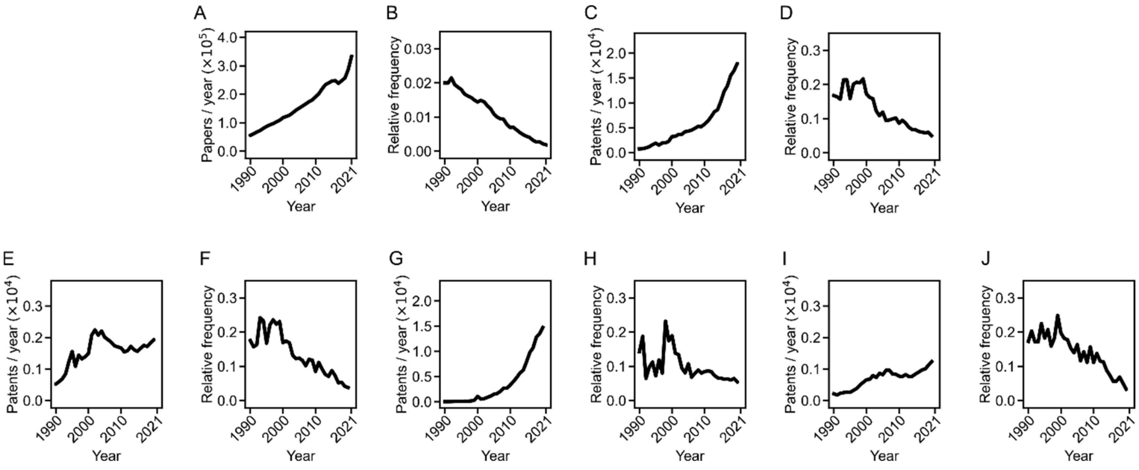


**Supplementary Fig. S3. New gene occurrence per paper or patent**. Related to Fig. 2A. **A**, **B**, The total annual counts of papers (**A**) and their proportion with new genes (**B**). **C**, **D**, The total annual counts of patents (**C**) and their proportion with new genes (**D**). **E**, **F**, The total annual counts of only US patents (**E**) and their proportion with new genes (**F**). **G**, **H**, The total annual counts of only Chinese patents (**G**) and their proportion with new genes (**H**). **I**, **J**, The total annual counts of only European patents (**I**) and their proportion with new genes (**J**).

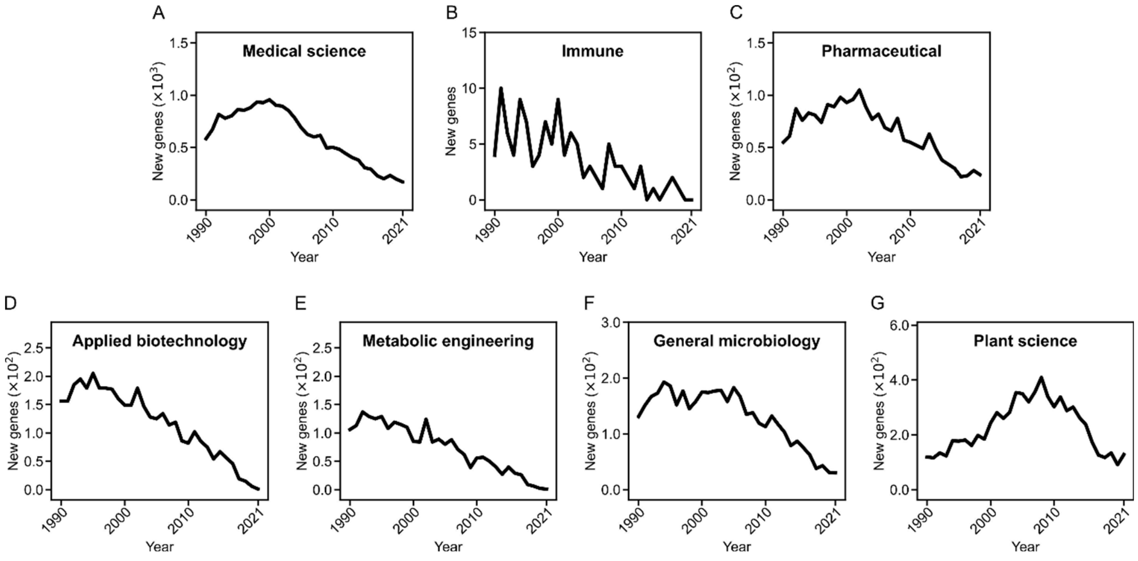


**Supplementary Fig. S4. Research on new genes in each thematic gene category.** Related to Fig. 2B–D**. A**–**G**, Annual counts of newly researched genes in each thematic gene category.

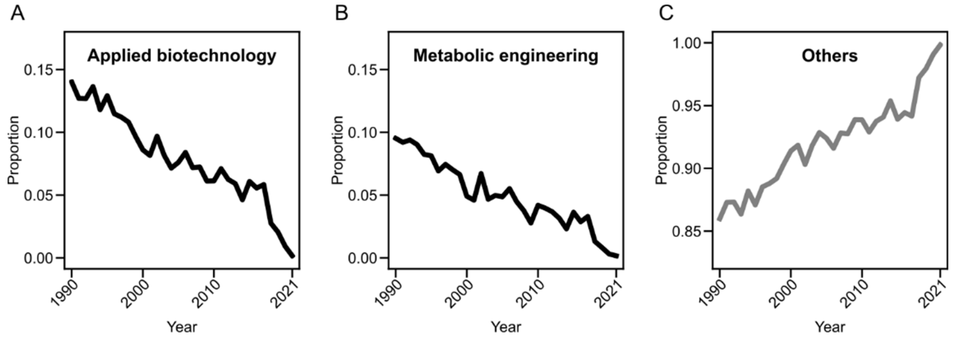


**Supplementary Fig. S5. Proportion of applied biotechnology or metabolic engineering genes among newly researched genes.** Related to Fig. 2C. The proportion each year (**A** and **B**) and that of the rest genes (**C**).

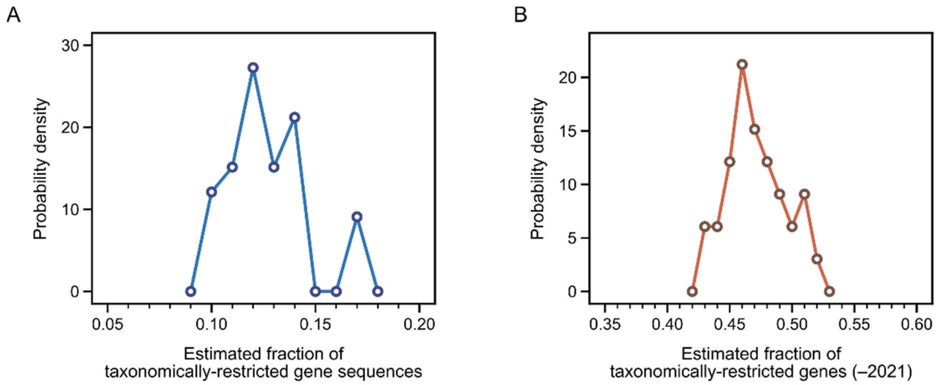


**Supplementary Fig. S6. Some estimates from our mathematical model.** Related to Fig. 2E. **A**, The probability distribution of the fraction of taxonomically-restricted gene (TRG) sequences among all the possible non-redundant sequences of research merit, estimated from our mathematical model (Sect. 4.7). It shows that TRGs makes up 9.9–17.3% of the sequences of research merit. This result is compatible with the existing knowledge that each sequenced genome contains 10–20% of genes as TRGs [40], as this known fraction is likely to set an upper bound of the TRGs of research merit given the TRGs' tendency of being less researched. **B**, The probability distribution of the fraction of TRGs among all researched genes until 2021, estimated from our mathematical model (Sect. 4.7). It shows that TRGs constitute 43.1–51.8% of the researched genes as of 2021. This result is compatible with the fraction (59.1%) of the genes present in single species (i.e., single entries) in UniProtKB/Swiss-Prot, because the fraction from UniProtKB/Swiss-Prot is possibly an overestimate with its missing species that may have the homologs of those single-entry genes.

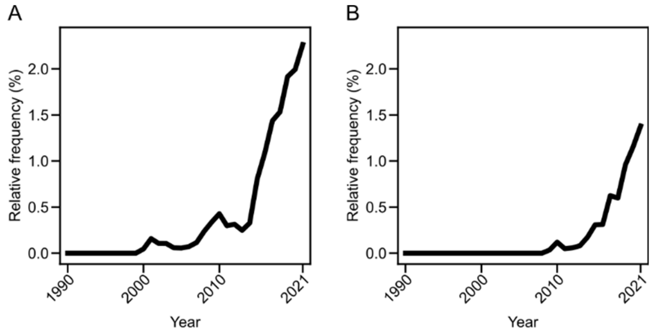


**Supplementary Fig. S7. Trend of third-generation sequencing.** The proportion of papers and patents with third-generation sequencing, among all the sequencing-related annual papers and patents (**A**) or only those for specific genes (**B**). Extrapolation of the data indicates 2 – 4 % adoption of this technology as of 2026.

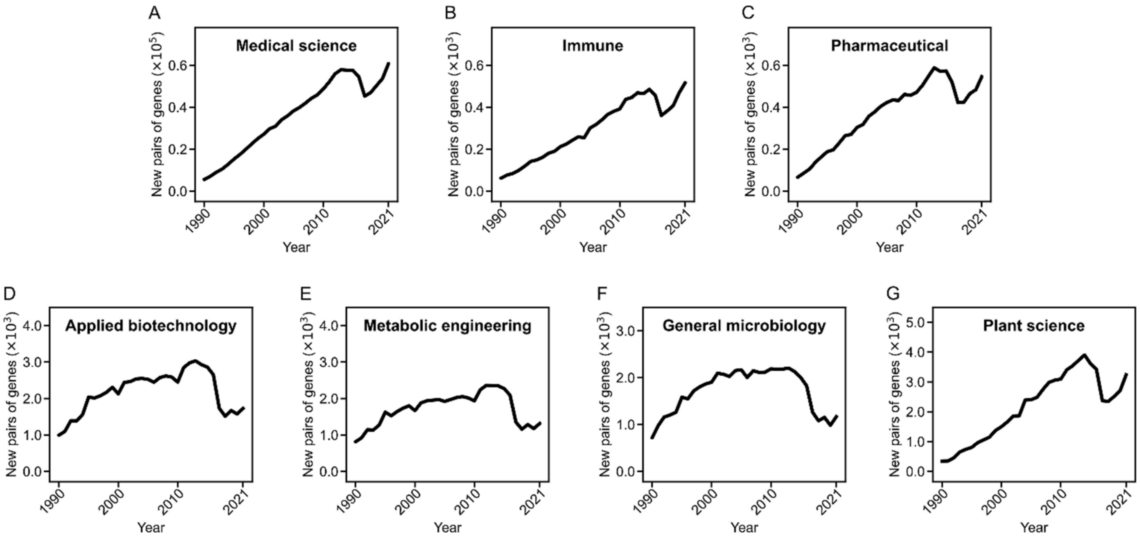


**Supplementary Fig. S8. Novel gene combinations in each thematic gene category.** Related to Fig. 3B–D. **A**–**G**, Annual counts of new pairs of genes in each thematic gene category.

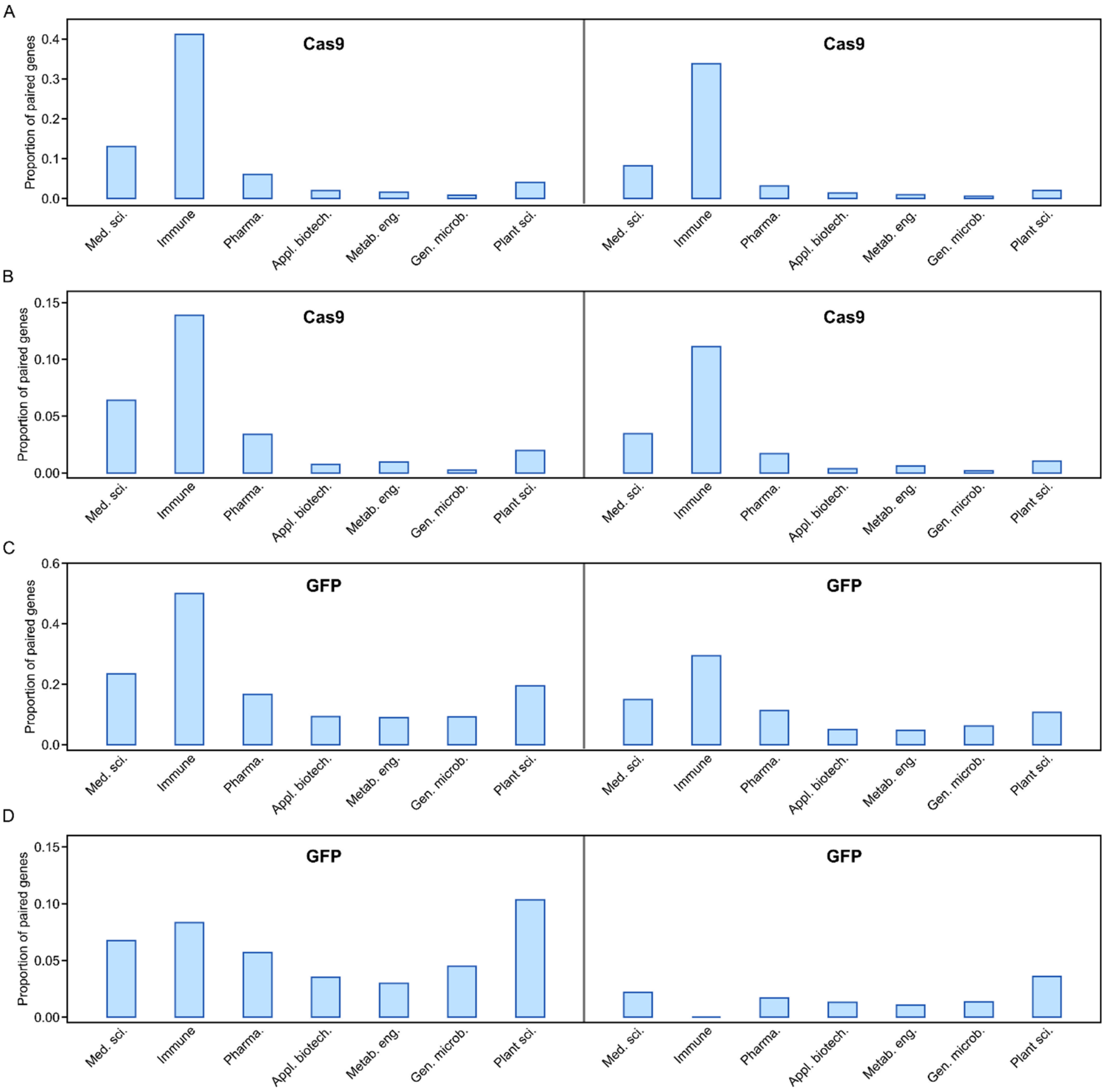


**Supplementary Fig. S9. Teaming with Cas9 or GFP in each thematic gene category.** Related to Fig. 3I. The proportion of genes in each thematic gene category, paired with Cas9 (**A** and **B**) or GFP (**C** and **D**) until its peak time (left) or earlier midpoint (right). For the definitions of the peak time and midpoint, refer to Sect. 4.9. In this teaming, we considered genes that debuted in papers/patents in 1990–2000 (**A** and **C**) and 2001–2011 (**B** and **D**), separately. Abbreviations: med. sci., medical science; pharma., pharmaceutical; appl. biotech., applied biotechnology; metab. eng., metabolic engineering; gen. microb., general microbiology; plant sci., plant science.

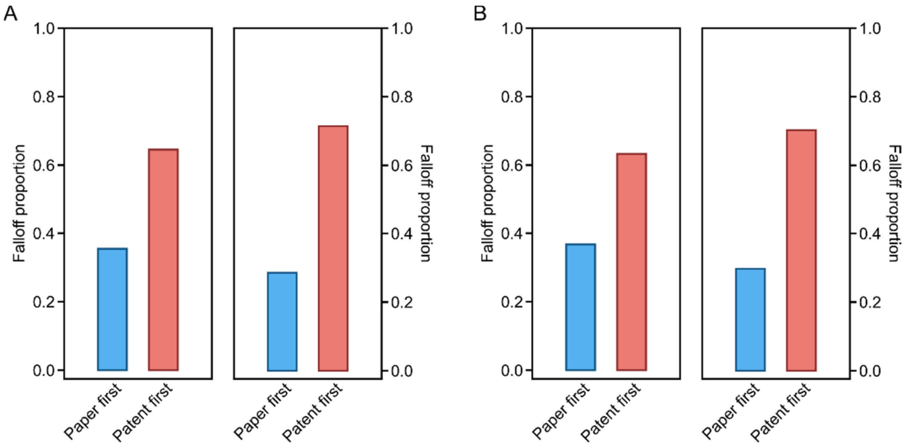


**Supplementary Fig. S10. Paper–patent falloff relationship with or without highly similar prior works.** Related to Fig. 4B. **A**, **B**, The proportion of genes whose paper falloffs precede the patent falloffs (blue) and the proportion of genes whose patent falloffs precede the paper falloffs (red). In this analysis, we first measured the content similarity of every pair of the papers (or patents) with each gene, using PaECTER [98]. If the total number of the papers (or patents) exceeds 5,000 for that gene, we only used 5,000 randomly selected from them to ensure computational feasibility. **A** shows the results with all these papers and patents, and **B** shows the results after filtering out the papers and patents highly overlapping with their prior works (Sect. 4.10). In **A** and **B**, we considered genes that debuted in papers/patents in 1990–2000 (left) and 2001–2011 (right), separately. For a fair comparison between the paper and patent falloffs, we set the endpoint of the paper time-series to the same year (2020) as the patent time-series.

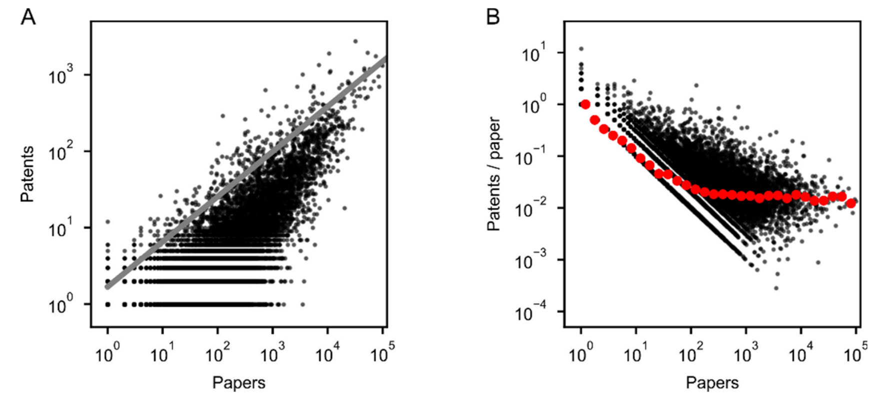


**Supplementary Fig. S11. Paper versus patent volumes.** Related to Fig. 4B. **A**, A scatter plot between the total paper and patent numbers of each gene. A gray straight line indicates $y = \alpha x^{\gamma}$ where $x$ and $y$ are the paper and patent numbers of a gene, respectively, $\gamma = 0.59$, and $\alpha = 1.68$. Most genes (98.8%) are placed below or on this line ($P < 10^{-4}$ and Sect. 4.11). In other words, the paper number of each gene tends to put an *upper limit* or a cap on its patent number. **B**, A scatter plot between the total paper number and patent-to-paper ratio of each gene (the median in each interval of the paper number is plotted in red). The more papers published for a gene, the fewer patents tend to be applied per paper, eventually at a level of ~1.6 patents/100 papers. In **A**, **B**, papers and patents are counted over the same periods (1990–2020) for their fair comparison. Only the genes with non-zero hits in both papers and patents are considered here.

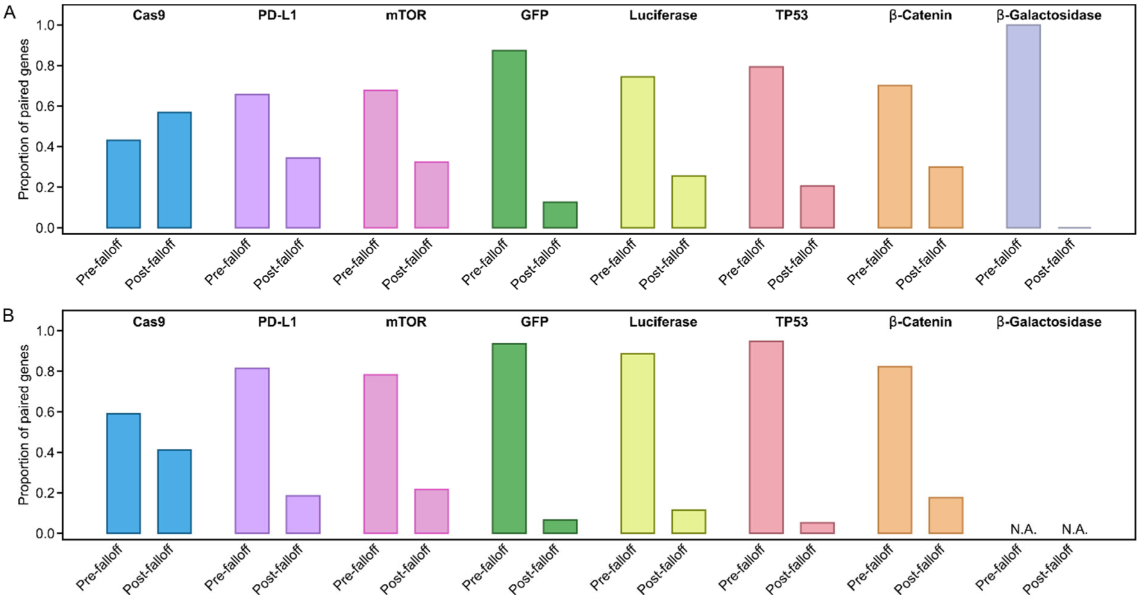


**Supplementary Fig. S12. Patent falloff and teaming with new iconic genes.** Related to Fig. 5B. **A**, **B**, The proportion of genes initially paired with a given iconic gene before ("Pre-falloff") or after/at ("Post-falloff") their patent falloffs. We here focus on the teaming until the peak time of this iconic gene (**A**) or the earlier midpoint (**B**) (Sect. 4.15). In the case of too small a number of genes itself whether paired with the iconic gene before or after the patent falloffs, this case is marked with "N.A." All the results were obtained with genes that debuted in papers/patents in a similar period (1990–2000). Genes with post-2000 debuts were not considered due to the fundamentally low overlap of their research periods with the early stages of the iconic genes.

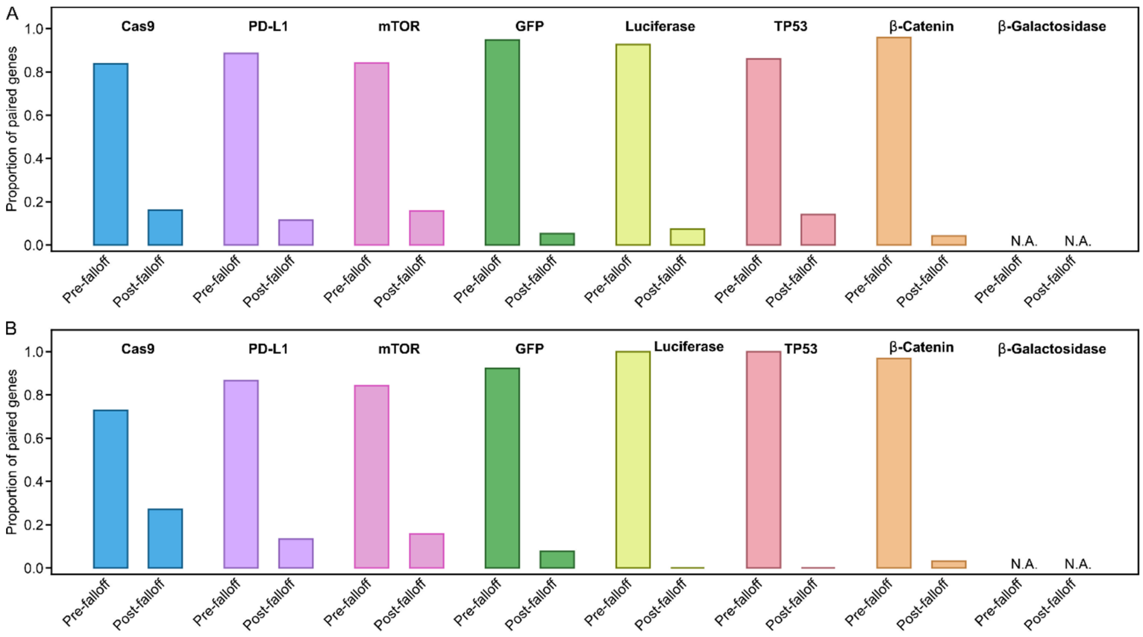


**Supplementary Fig. S13. Paper publications and teaming with new iconic genes.** Related to Fig. 5C. **A**, **B**, The proportion of genes initially paired with a given iconic gene before ("Pre-falloff") or after/at ("Post-falloff") their paper falloffs, among the genes with this teaming after/at the patent falloffs. We here focus on the teaming until the peak time of this iconic gene (**A**) or the earlier midpoint (**B**) (Sect. 4.15). In the case of too small a number of genes itself whether paired with the iconic gene before or after the paper falloffs, this case is marked with "N.A." All the results were obtained with genes that debuted in papers/patents in a similar period (1990–2000). Genes with post-2000 debuts were not considered due to the fundamentally low overlap of their research periods with the early stages of the iconic genes.

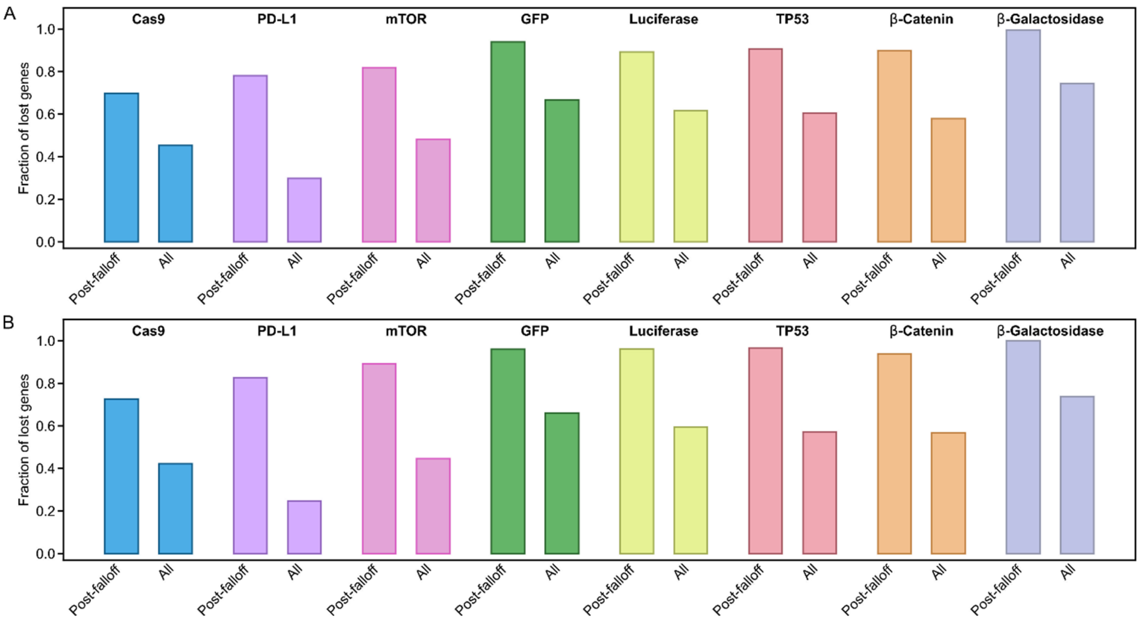


**Supplementary Fig. S14. Missing opportunities for the teaming with iconic genes.** Related to Fig. 5D. **A**, **B**, The estimated proportion of genes that would lose the teaming with a given iconic gene, among the genes that initially teamed with this iconic gene with their limited patent volumes ("Post-falloff") or among all the genes that teamed with this iconic gene ("All"). We here focus on the initial teaming until the peak time of that iconic gene (**A**) or the earlier midpoint (**B**) (Sect. 4.16). The results were obtained with genes that debuted in papers/patents in a similar period (1990–2000). Genes with post-2000 debuts were not considered due to the fundamentally low overlap of their research periods with the early stages of the iconic genes.

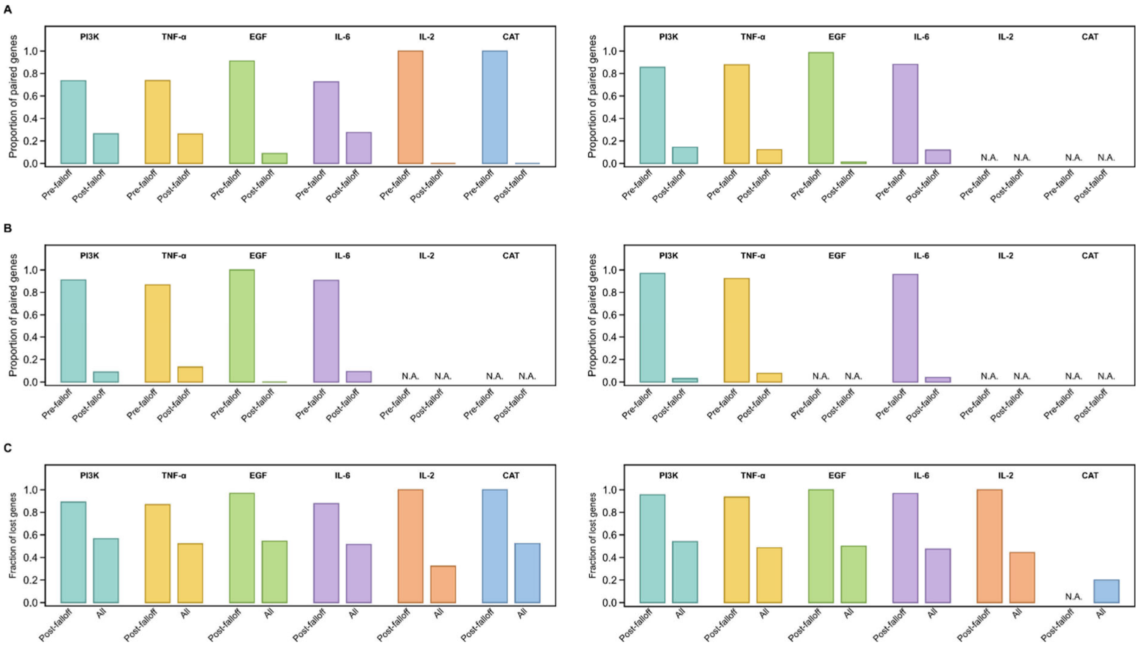


**Supplementary Fig. S15. Initial teaming with the genes in Supplementary Table S1, excluding iconic genes.** Related to Fig. 5B–D. **A**, The proportion of genes initially paired with a given gene in Supplementary Table S1 before ("Pre-falloff") or after/at ("Post-falloff") their patent falloffs. **B**, The proportion of genes initially paired with a given gene in Supplementary Table S1 before ("Pre-falloff") or after/at ("Post-falloff") their paper falloffs, among the genes with this teaming after/at the patent falloffs. **C**, The estimated proportion of genes that would lose the teaming with a given gene in Supplementary Table S1, among the genes that initially teamed with this gene with their limited patent volumes ("Post-falloff") or among all the genes that teamed with this gene ("All"). In **A**–**C**, we focus on the initial teaming until the peak time of that gene (left) or the earlier midpoint (right) (Sect. 4.15, 4.16). In the case of too small a number of genes itself for the relevant teaming, this case is marked with "N.A." All the results were obtained with genes that debuted in papers/patents in a similar period (1990–2000). Genes with post-2000 debuts were not considered due to the fundamentally low overlap of their research periods with the early stages of those genes in Supplementary Table S1.

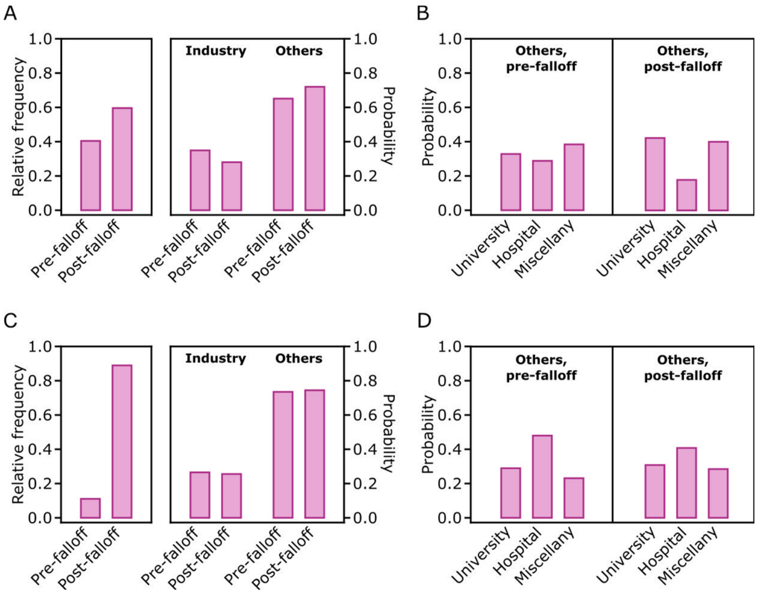


**Supplementary Fig. S16. Clinical trial analysis.** **A**, Adopted from Fig. 4C based on the trials and genes in papers. **B**, The probability of university, hospital, or other non-industry sponsorship for a clinical trial with each gene in a given case ("Others, pre-falloff" or "Others, post-falloff") from the right side of **A** (Sect. 4.13). **C**, **D**, Instead of considering clinical-trial papers as **A**, **B**, we directly screened the XML file of ClinicalTrials.gov for trial-related genes and data (Supplementary Table S4), and repeated the analysis in **A** (**C**) or **B** (**D**). In **A**–**D**, we only analyzed the clinical-trial genes that debuted in papers/patents in 2002–2011, regarding the first year of the mandated trial posting to ClinicalTrials.gov (Sect. 4.13).

| Year / Rank | 1990–1994 | 1995–1999 | 2000–2004 | 2005–2009 | 2010–2014 | 2015–2019 | 2020 | 2021 |
|---|---|---|---|---|---|---|---|---|
| 1 | β-Gal | GFP | GFP | GFP | GFP | Cas9 | Cas9 | Cas9 |
| 2 | IL-2 | β-Gal | Luc | TP53 | Luc | Luc | Luc | Luc |
| 3 | IL-6 | Luc | TP53 | Luc | TP53 | mTOR | mTOR | PD-L1 |
| 4 | CAT | TP53 | β-Gal | PI3-K | PI3-K | PI3-K | PI3-K | mTOR |
| 5 | EGF | TNF-α | PI3-K | β-Catenin | mTOR | β-Catenin | IL-6 | β-Catenin |

**Supplementary Table S1. Frequent genes in novel gene pairs in each period.** Five genes of the largest shares in new pairs of genes in papers or patents in each 5-year period up to 2019, and in 2020 and 2021 separately. Abbreviations: β-Gal, β-galactosidase; IL-2, interleukin-2; IL-6, interleukin-6; CAT, chloramphenicol acetyltransferase; EGF, epidermal growth factor; GFP, green fluorescent protein; Luc, luciferase; TP53, tumor suppressor protein p53; TNF-α, tumor necrosis factor-α; PI3-K, phosphatidylinositol 3-kinase; mTOR, mammalian target of rapamycin; Cas9, CRISPR associated protein 9; PD-L1, programmed cell death ligand 1.

| Section | Attribute | Remark |
|---|---|---|
| Protein description | Recommended name | |
| | Alternative name | |
| | Allergen | |
| | CD antigen | |
| | INN (International Nonproprietary Names) | |
| Gene names | Primary name | |
| | Synonyms | |
| | Ordered locus number | Excluded from our analysis |
| | ORF (open reading frame) number | Excluded from our analysis |

**Supplementary Table S2. Available attributes of gene and protein names in UniProtKB/Swiss-Prot entries.** Both **"**Ordered locus numbers" and "ORF numbers" were excluded in the listing of our gene names, regarding their frequent use in the cases of not-fully-identified gene products. All the other attributes were considered for the listing of the gene names.

| | | Our algorithm | | PubTator3 | |
|---|---|---|---|---|---|
| Attribute | Test group | Precision | Recall | Precision | Recall |
| Debut year | 1990–2000 | 0.87 | 0.69 | 0.95 | 0.69 |
| Debut year | 2001–2011 | 0.93 | 0.71 | 0.93 | 0.62 |
| Paper & patent counts | Top 10% | 0.87 | 0.66 | 0.92 | 0.58 |
| Paper & patent counts | Top 10–50% | 0.91 | 0.74 | 0.95 | 0.55 |
| Paper & patent counts | Lower 50% | 0.90 | 0.74 | 0.92 | 0.46 |
| Patenting region | US | 0.91 | 0.71 | N.A. | N.A. |
| Patenting region | China | 0.86 | 0.76 | N.A. | N.A. |
| Patenting region | Europe | 0.89 | 0.73 | N.A. | N.A. |
| Thematic category | Medical science | 0.89 | 0.75 | 0.91 | 0.64 |
| Thematic category | Applied biotechnology | 0.92 | 0.67 | 0.93 | 0.21 |
| Thematic category | General microbiology | 0.93 | 0.72 | 0.98 | 0.18 |
| Thematic category | Plant science | 0.90 | 0.72 | 0.93 | 0.48 |
| Minimum | | 0.86 | 0.66 | 0.91 | 0.18 |
| Maximum | | 0.93 | 0.76 | 0.98 | 0.69 |
| Mean | | 0.90 | 0.72 | 0.94 | 0.49 |

**Supplementary Table S3. Accuracy of gene search.** The precision and recall of our gene search algorithm for papers and patents are presented across different test groups of the attributes of query genes (Sect. 4.3). These attributes include the debut year, the total paper and patent counts, the patenting region (at least once), and the thematic gene category. For each test group, 100 papers and patents were examined in a nearly 1:1 ratio, except for the patenting-region case where only 100 patents from a given region were used. For comparison, the precision and recall of PubTator3 are also presented, based on the same paper sets of the test groups. Because PubTator3 does not cover patents, only the papers were used and the patenting-region cases are marked with “N.A.”

| XML field | Utility in our analysis |
|---|---|
| NCT number | Trial identifier |
| Brief title | Gene search |
| Official title | Gene search |
| Brief summary | Gene search |
| Detailed description | Gene search |
| Conditions | Gene search |
| Keywords | Gene search |
| Intervention names | Gene search |
| Intervention descriptions | Gene search |
| Primary outcome measures | Gene search |
| Primary outcome descriptions | Gene search |
| Secondary outcome measures | Gene search |
| Secondary outcome descriptions | Gene search |
| Eligibility criteria | Gene search |
| Start date | Trial year |
| Study first posted | Trial year if "Start date" is unavailable |
| Lead sponsor name | Sponsor classification |
| Lead sponsor class | Sponsor classification |
| Collaborator names | Sponsor classification |
| Collaborator classes | Sponsor classification |
| Last update posted | |
| Overall status | |
| Phase | |
| Study type | |
| Study first submitted | |

**Supplementary Table S4. ClinicalTrials.gov XML fields considered for our clinical-trial analysis**. Descriptive fields were examined to identify trial-related genes by the string-matching algorithm in Sect. 4.3. The NCT number was used as the trial identifier. Other fields were extracted to determine the trial year and sponsor type.
